# Benchmarking of PROTAC docking and virtual screening tools

**DOI:** 10.1101/2023.08.30.555318

**Authors:** Evianne Rovers, Matthieu Schapira

## Abstract

Proteolysis targeting chimeras (PROTACs) are bifunctional compounds that recruit an E3 ligase to a target protein to induce ubiquitination and degradation of the target and are pioneer molecules in the field of proximity pharmacology. Rational PROTAC design is a challenging process and novel computational tools have emerged that attempt to predict the ternary complexes created by PROTACs and identify PROTAC candidates. To compare the performance of recent PROTAC design and screening methods, a benchmark was developed to test the ability of these tools to 1) predict the ternary complexes observed in crystal structures and 2) dissociate active from inactive PROTACs. Unlike traditional protein-protein complex prediction software, the PROTAC virtual screening methods often generate successfully PROTAC-induced protein complex structures observed crystallographically, but these experimentally validated predictions are not dissociated from dozens or more of other predicted structures. PROTAC virtual screening efficiency is unclear and highly variable, in part due to the limited size of experimental datasets and the low number of negative controls. Defining ubiquitination zones within cullin-RING complexes does not improve predictions, but conformational arrangements can sometimes be found that are exclusively associated with active PROTACs. Computer assisted PROTAC design is still in its infancy. Pioneering tools highlight the promises and challenges in the field and may be more valuable when guided by clear structural and biophysical data and validated on specific chemical series.

## Introduction

Proximity induced pharmacology represents a new paradigm in drug discovery where a molecule is designed to stabilize a specific and non-natural interaction between two proteins leading to a desired phenotype^1–3^. Among these types of molecules, proteolysis targeting chimeras (PROTACs) are closest to proof-of-concept, with multiple drugs in clinical trials^1^. PROTACs are heterobifunctional compounds where small molecule ligands for an E3 ubiquitin ligase and a protein target are chemically linked, thereby inducing the ubiquitination and subsequent degradation of the target. Unlike occupancy-based drugs that typically inhibit their targets stoichiometrically, PROTACs catalyze the degradation of their target, a sub-stoichiometric mechanism allowing lower dosage; they can also exploit non-functional binding sites of otherwise untractable targets and are active only in tissues expressing the recruited E3 ligase, which can lead to reduced toxicity^4^.

The discovery of a PROTAC can be a frustrating trial-and-error process where a multitude of linkers and attachment points are tested with ligands for different E3 ligases in the hope that one combination produces an active molecule. The optimization of active PROTACs typically consists of a medicinal chemistry exploration focused on cell permeability and potency that can be systematic and highly customized^5,6^, but is rarely structure-enabled.

Given the significant impact of computer-assisted drug design (CADD) in hit optimization and, to a lesser extent, hit discovery for traditional occupancy-based molecules, a reasonable question is whether CADD can also make significant contributions in the discovery and optimization of PROTACs. On the one hand, PROTAC virtual screening appears as a daunting task considering that (1) the exact binding pocket, at the interface of the two proteins, is not defined a priori, (2) the ab initio docking and scoring of a small molecule to a binding site captured in its apo state is unreliable and (3) CADD typically underperforms with large and flexible molecules. On the other hand, predicting the structure of a ternary complex (E3 ligase-PROTAC-target protein) may be seen as a simplified protein-protein docking exercise, where (1) the available conformational space is drastically constrained by the PROTAC, (2) the docking poses of the PROTAC chemical handles (i.e. ligands occupying the E3 and the target) are already known and (3) the complexity of the system may be reduced to the flexibility of the linker.

New computational methods, protocols and tools are emerging to predict ternary PROTAC complex structures and screen virtually libraries of PROTAC candidates. Results reported by the developers seem encouraging in some cases but cannot be compared when focused on different case studies. Additionally, ternary structures are sometimes “predicted” by docking monomeric structures themselves extracted from the crystallized ternary complex, which does not reflect the challenges of docking two proteins that never met.

To better appreciate the current value of CADD for PROTAC discovery, we assembled a benchmark of 10 experimental ternary complex structures including three E3 ligases to test the ability of three recently reported computational tools to predict these structures. We also tested whether these tools could dissociate active from inactive PROTACs in 5 PROTAC discovery campaigns reported in the literature. We find that experimental protein-protein interfaces are reproduced with sometimes high accuracy, which may enable linker optimization, but these high-accuracy structures are computationally ranked lower than other more dissimilar ones. We also discuss the biological relevance of crystallized ternary complexes. The efficiency of PROTAC virtual screening is not clear in our hands, even when incorporating geometrical constraints related to the orientation of the substrate protein in the context of the cullin-ring multi-protein complex. We do observe that in some cases, active PROTACs are enriched in a specific conformational arrangement of the protein-protein complex, which could be exploited for lead optimization.

## Benchmark and tools

A few pioneer computational tools have recently been reported, and all rely on the prediction of a ternary complex structure^7–9^, the assumption being that PROTACs that are compatible with the formation of a stable complex are more likely to be active.

PRosettaC^7^ and MOE^8,9^ predict multiple putative protein-protein complex structures where each protein is bound to a PROTAC chemical handle (available in the PDB), but without the PROTAC linker. The conformational space of the PROTAC is sampled independently, in the absence of protein for MOE and in the protein complex for PRosettaC. PROTAC conformations compatible with protein complex structures are then identified and clustered. Cluster sizes are used to rank ternary complexes. Conversely, ICM (Molsoft, San Diego) continuously minimizes the energy of the ternary complex with the flexible PROTAC and uses the predicted energy of the complex for ranking (Fig. 1).

**Figure 1.**
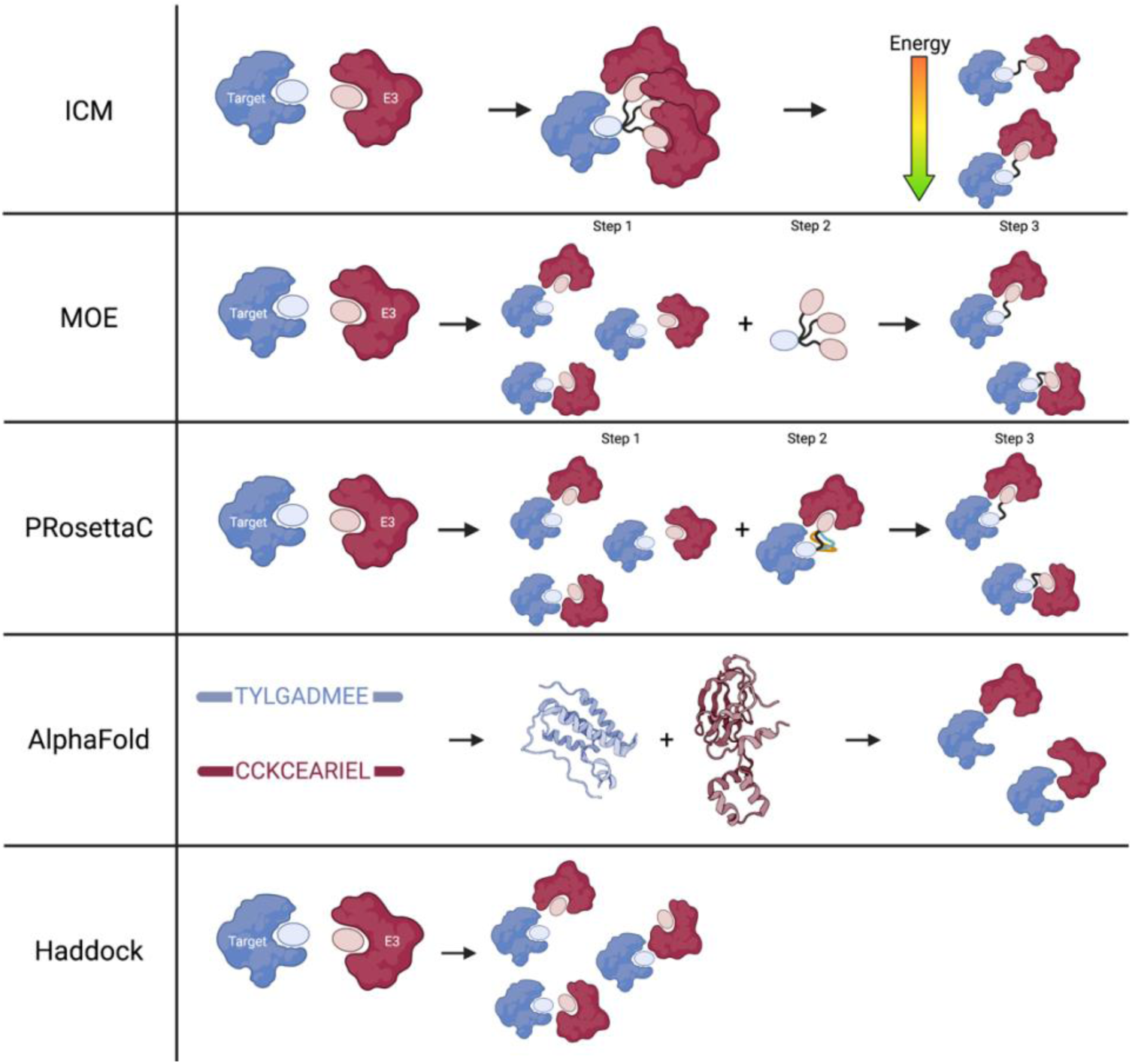
Complex prediction using different methodologies (AlphaFold, Haddock, MOE, Molsoft and PRosettaC). ICM assembles the ternary complex and runs MonteCarlo simulation with flexible linker and residues surrounding the PROTAC to obtain multiple ternary complex conformations ranked based on energy. MOE and PRosettaC perform protein-protein docking followed by conformational search of the PROTAC. Ranking is based on cluster population. AlphaFold uses machine learning algorithm to predict monomer structures and assemble protein complexes. Haddock predicts binary complex using provided interaction residue information.

While a few prospective or retrospective studies have been reported for some of these computational tools, they focus on different targets and PROTACs and cannot be compared. Here, we systematically benchmark PRosettaC, MOE and ICM for their ability to predict ternary complex structures, and to dissociate active from inactive PROTACs. We also test two protein-protein interaction prediction tools that ignore the PROTAC for their ability to predict binary protein complex structures observed in experimental PROTAC ternary complexes: (1) HADDOCK^10^ which docks proteins bound to their chemical handles with defined distance restraints between residues; (2) AlphaFold-multimer^11^, that requires as input two protein sequences and uses deep learning to build a binary complex structure (Fig. 1). We provide all data for this benchmarking exercise in the hope that it will be used and augmented by others as more PROTACs and inactive analogs are reported, more ternary complex structures are deposited in the Protein Data Bank (PDB), and more and improved computational methods are developed to rationally design PROTACs.

## Methods

### Replication of previous results

To replicate the results published with MOE^8,9^ and PRosettaC^7^, two test cases were used: 6BOY (CRBN-BRD4BD1)^12^ and 5T35 (VHL-BRD4BD2)^13^. These test cases were the first crystal structures of a ternary complex (E3-PROTAC-Target) to be published in the PDB and were used by both developers. In both cases, the monomeric structure of each protein used for docking was extracted from the crystal structure of the ternary complex. For PRosettaC, the input files are available online^7^ and for MOE, the input files were generated by separating the crystal structure of the complex into the proteins and their chemical handle. Both methods were run using the default settings. The root mean square deviation of the backbone (Ca-RMSD) of the target protein is calculated after superimposing the E3 ligase from the predicted and crystallized complex. A detailed description of the protocol is provided in a separate report^14^.

### Accuracy prediction of ternary complex

To evaluate the ability of the complex prediction methods to accurately predict the ternary complex, we chose ten test cases published in the PDB (Tab. 1). It includes a variety of E3 ligases (CRBN, VHL, cIAP) and target proteins (kinases, bromodomains, SMARCA2/4 and WDR5). The VHL-WDR5 has two complexes available in the PDB bound to PROTACs with similar handles and different linker length. The PROTACs induce two different conformations, which illustrates the importance of the linker in the ternary complex prediction. To prepare the input files for all methods, a crystal structure of the unbound protein in complex with its chemical handle was used. Definitions of chemical handles and linkers are described in table S2. The 3D structures were extracted from the PDB, other chains and ligands were deleted, and the biologically relevant oligomeric state was generated with ICM. In case bonds and atoms present in the PROTAC were missing in the chemical handle, the chemical handle was adjusted to represent the chemical handle in the PROTAC. For Haddock, PRosettaC and MOE, the PDB files were saved and for Molsoft the icm-object files. For MOE, the PDB files were opened and the ‘QuickPrep’ module was run and the proteins saved as MOE object files. The PROTAC was saved as either a SDF file (Molsoft), a MOE database (MOE) or a smiles.txt file (PRosettaC). For AlphaFold, sequences were cut to represent the apo-crystal structure and saved in a fasta file. To guide protein-protein docking in Haddock, residue restraints were chosen by selecting residues that are <5Å distant from the chemical handle and are solvent accessible (Tab. S1). All input files are available on https://zenodo.org/record/8298749. Each methodology was run using the default settings, except Haddock for which the number of output structures for the first stage was increased to 10000 and structures for analysis to 400. Analysis is based on 1) the Ca-RMSD (Å) of target protein 2) the RMSD of the interface (<10Å residues in target protein from E3 ligase), 3) PROTAC RMSD, 4) target chemical handle RMSD, 5) the ranking of the near-native complex and 6) the Ca-RMSD (Å) of the top-ranking complex.

**Table 1.**
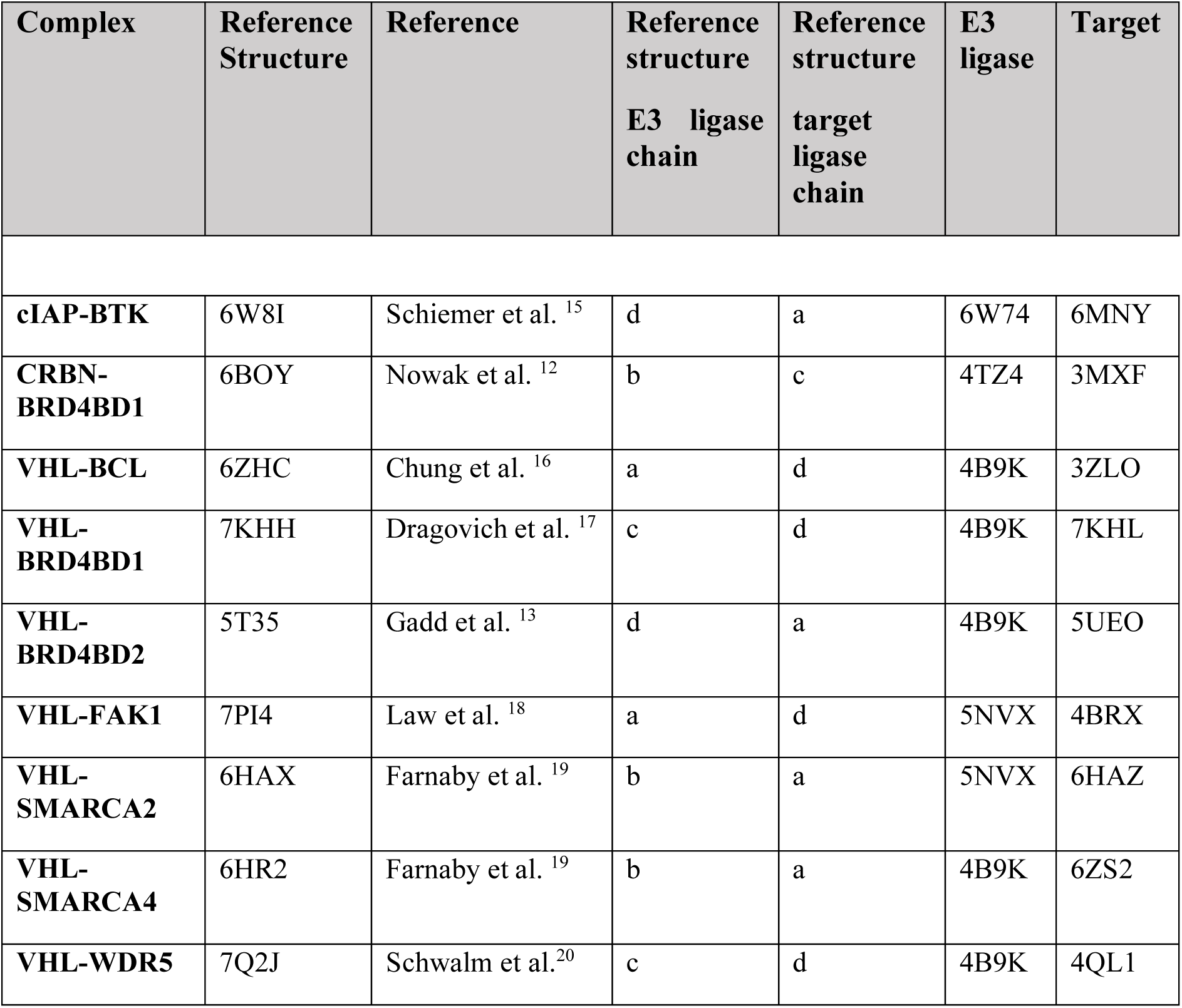
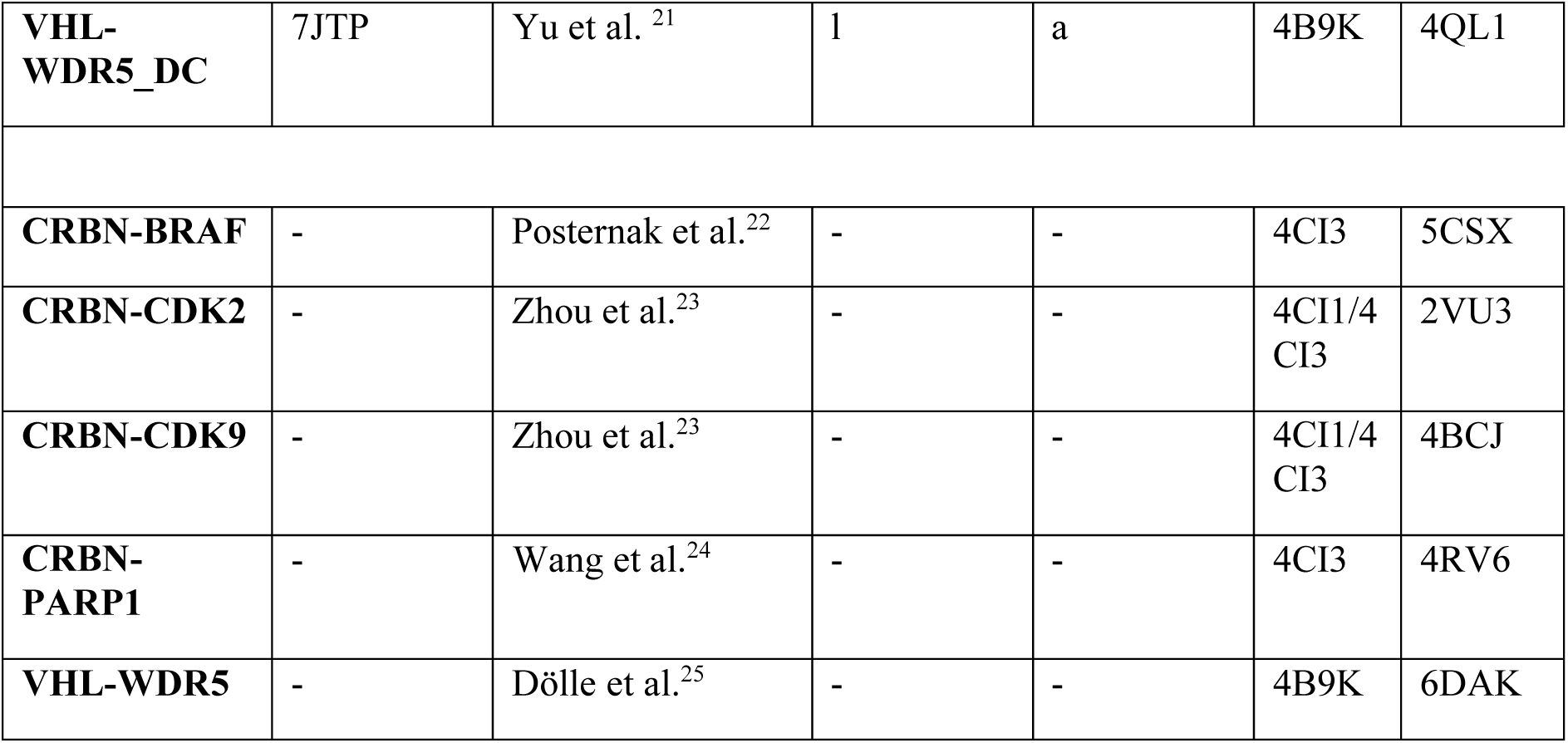
Benchmark data set. Top: PDB code of experimental ternary complex and isolated protein structures used to evaluate the accuracy of the docking step. Bottom: PDB codes of protein structures used to evaluate the accuracy of PROTAC virtual screening.

### Distinguishing between active and non-active PROTACs

To evaluate the accuracy in differentiating active from non-active PROTACs, five test cases studies were chosen CRBN-BRAF^22^, CRBN-CDK2^23^, CRBN-CDK9^23^, CRBN-PARP1^24^ and VHL-WDR5^25^. All test studies have active and non-active PROTACs with ratio of actives/total compounds of 5/8 (62.5%%), 6/20 (30%), 8/20 (40%), 8/13 (62%%) and 6/16 (37.5%) respectively (Tab. S3-8). To prepare the input files for all methods, unbound protein-structures in complex with their chemical handle was taken from the the PDB and prepared as described previously for MOE, Molsoft and PRosettaC (Tab. 1). Each methodology was run using the default settings. For MOE and PRosettaC, PROTAC ranking is determined by the cluster population sizes. Since Molsoft does not include ranking of the PROTACs, a similar ranking protocol as MOE and PRosettaC is performed by clustering the ternary complexes using the clustering option in ICM and ranking the PROTACs based on cluster population size.

### Searching for optimal ubiquitination zones

Another metric to determine the relevance of the ternary complex conformation is to measure the distance between lysine residues in the target and the E2 ligase in the RING complex after superimposition. Presence of a lysine in the ubiquitination zone could indicate a viable ternary complex conformation for ubiquitin transfer^26^. The PhosphositePlus database provides the post-translational modification status of lysine residues in the target^27^. Both the ubiquitination status and acetylation status were considered, since acetylation can precede ubiquitination^28^. All crystal structures containing a PROTAC ligand for VHL were downloaded from the PDB. The ternary complexes were superimposed on the cullin system (PDB:5N4W)^29^. Since 5N4W does not include the structure of the E2 ligase nor ubiquitin, the minimum distance was calculated between target lysine residues (known to be ubiquitinated or acetylated) and the adapter protein RBX1 to which the E2 ligase binds.

### Looking for conformations that dissociate active from inactive PROTACs

We looked at conformational clusters that are highly enriched in active PROTACs vs inactive PROTACs. For each test case, the conformational cluster centres were extracted from the PROTAC virtual screening produced by ICM. All the cluster centres for all the ligands were grouped into superclusters with cut-off of 4 Å. We looked at the number of actives and inactive PROTACs per supercluster. For the VHL-WDR5, the “actives-only” supercluster was compared to the crystal structures of the VHL-WDR5 complex (PDB: 7JTP^21^,7Q2J^30^) and Ca-RMSD calculated.

### Virtual screening using a ternary complex crystal structure

To simulate a PROTAC optimization study, we extracted the protein-protein complexes of the VHL-WDR5 crystal structures (PDB: 7JTP^21^,7Q2J^30^). The chemical handles were adjusted depending on the PROTACs library screened. We created a MOE database of the protein-protein complexes and ran the MOE protocol with the option for using pre-generated protein-protein complexes using the PROTAC library for VHL-WDR5. The enrichment of active PROTACs in the top, top 3 or top 5 selections was calculated to determine the efficacy of PROTAC virtual screening using crystal structures as input.

## Results

The benchmarking exercise was composed or four steps: we first verified that we could replicate previously published results reported by the developers of the MOE protocol and PRosettaC. We next evaluated the accuracy of ternary complex prediction and the ability to distinguish active from inactive PROTACs. Finally, we evaluated strategies to increase virtual screening efficiency by defining optimal ubiquitination zones, identifying conformational clusters enriched in active molecules or using crystal structures for PROTAC virtual screening.

### Ternary complex prediction

#### Benchmark data set

We assembled a benchmark data set composed of 10 test cases with one CRBN mediated complex, one complex with the cIAP ligase, and eight containing ligase VHL (Tab. 1). Two crystal structures of the VHL-WDR5 complex (PDB: 7Q2J^30^ and 7JTP^21^) are induced by different PROTACs and the WDR5 protein adopts a different orientation with each PROTAC.

#### Replication of previous results

Re-docking protein monomers extracted from the crystal structure of the ternary complex produced results similar to those previously published by developers using both MOE and PRosettaC (Tab. 2). The predicted CRBN-BRD4BD1 complex (later referred to as near-native complex) closest to the crystal structure (6BOY) had a carbon-α (“Ca” i.e. protein backbone) RMSD of 1.68Å and 3.21Å using PRosettaC which is close to the published Ca-RMSD of 2.26Å. The VHL-BRD4BD2 crystal structure (5T35) was predicted with an accuracy of 1.79Å and 4.09Å Ca-RMSD with PRosettaC compared to the published value of 2.26Å. The variation in Ca-RMSD and ranking between the runs and the predicted result can be attributed to the low reproducibility within the protocol which is also observed in a reproducibility study (Fig. S1). A large variety in Ca-RMSD and ranking is observed for PRosettaC when running the protocol three times using the same input files (Fig. S1).

**Table 2.** Replication of published ternary complex predictions using MOE^8,9^ and PRosettaC^7^. Each prediction was conducted twice independently with MOE and PRosettaC, on two ternary complexes, as published by the software developers. MOE: the number of near-native complexes/total structures when looking at Ca-RMSD (<10Å). The CAPRI metrics is high/medium/acceptable for interface RMSD (<1 Å / >1 Å & <2 Å / >2 Å & <4 Å). The percentages for both are the number of structures meeting the criteria / total structures. PRosettaC: The accuracy (Ca-RMSD) and rank of the most accurate prediction are provided. Ca-RMSD represents the backbone RMSD between the protein binding pose of the docked and crystallized complexes. A rank of x/y means that the most accurate protein binding pose was ranked x out of y predicted conformations. Ranking is based on the cluster size.

| Method | Test case | Metric | Published result | Replication result |  |
| --- | --- | --- | --- | --- | --- |
| MOE | CRBN-BRD4BD1 | Ca-RMSD | 4/1063 (0.4%) | 0/442 (0%) | 0/432 (0%) |
|  |  | CAPRI | 0/4/0 (0.4%) | 0/0/0 (0%) | 0/0/0 (0%) |
|  | VHL-BRD4BD2 | Ca-RMSD | 979/1692 (57.9%) | 256/356 (73%) | 207/289 (72%) |
|  |  | CAPRI | 50/950/198 (70.8%) | 40/0/104 (40%) | 40/0/94 (43%) |
| PROsettaC | CRBN-BRD4BD1 | Ca-RMSD | 2.26 Å | 1.68 Å | 3.21 Å |
|  |  | Ranking | 2/22 | 9/36 | 7/36 |
|  | VHL-BRD4BD2 | Ca-RMSD | 2.26 Å | 1.79 Å | 4.09 Å |
|  |  | Ranking | 2/46 | 18/114 | 2/110 |

Drummond et al. did not publish the Ca-RMSD per complex, but provided the ratio of near-native structures (<10Å RMSD) among the biggest conformational cluster. Additionally, the predictions were analyzed according to the CAPRI criterion that looks at the interface RMSD^31^. As published, we find high number of near-native structures for the VHL test case, but not for the CRBN test case. In fact, in our hands near-native CRBN complex structures were found in other clusters (30 near-native structures out of 3363 predicted conformations (0.9%)). Our results for the VHL complex are better than published (72% vs 57.9% near-native structures) which may reflect the fact that we use a more recent version of MOE.

### Accuracy prediction of near-native complex

Having established that we could reasonably reproduce previously published results that used monomeric protein structures extracted from crystallized ternary complexes, we next evaluated the accuracy of the docking models when using experimental structures of the unbound proteins, a task that better represents prospective discovery efforts. We first ran the docking simulations in triplicate to test the reproducibility of predictions on two test cases: CRBN-BRD4BD1 and VHL-FAK1. Overall, the results were consistent, both in terms of the accuracy and rank of the best (a.k.a. “near-native”) prediction, and in the accuracy of the top-ranking structure (Fig. S1). In particular, results produced with MOE were highly reproducible, while some variations were observed with ICM, especially in the accuracy of the top-ranking structure. We next ran the docking simulations on all ten complexes listed Table 1. ICM generally produced the most accurate structure, with an average Ca-RMSD of 5Å with the experimental structure, followed by MOE and PRosettaC with averages of 9Å and 11Å Ca-RMSD respectively (Fig. 2A). Haddock (18.6 Å) and AlphaFold (28.0 Å) did not perform as well, supporting the idea that addition of the PROTAC in the simulations imposes favorable conformational constraints on the protein-protein interface. Modelling of the interface between the two proteins is more important than the whole complex, since the protein-protein interactions are responsible for the stability of the complex and the PROTAC binds at the protein interface. When looking at the interface RMSD, the accuracy improves for all methods, with RMSD values below 10Å for most test cases. The PROTAC RMSD varies greatly across test cases and methods but was almost always above 2Å, which would be a minimum accuracy necessary to rationally design improved analogs. Similarly, the relative positioning of the chemical handles was often above 4 Å, and therefore poorly inform minimal linker length. Simulations with PRosettaC failed to produce a ternary complex for cIAP-BTK and VHL-BCL, even after adjusting chemical handles or using different crystal structures for both the target and E3 ligase. Striking differences in computing time distinguished the benchmarked tools. ICM and MOE used on average 14 and 45 CPU hours respectively while AlphaFold took 193 and PRosettaC 431 CPU hours (Fig. 2B and Tab. 3). Haddock was run on a webserver (https://wenmr.science.uu.nl/haddock2.4/) and was consistently the fastest. Importantly, all tested tools predicted multiple structures, and the more output structures generated, the more likely an accurate near-native solution was found (Fig. 2C). For example, an accurate conformation was more often produced by MOE and ICM but was one among 3637 and 1348 predicted structures on average respectively. This contrasts strikingly with the 10, 25 and 250 structures predicted by Haddock, AlphaFold and PRosettaC. A critical question is then whether the near-native structure was one of the top-ranking predictions.

**Figure 2.**
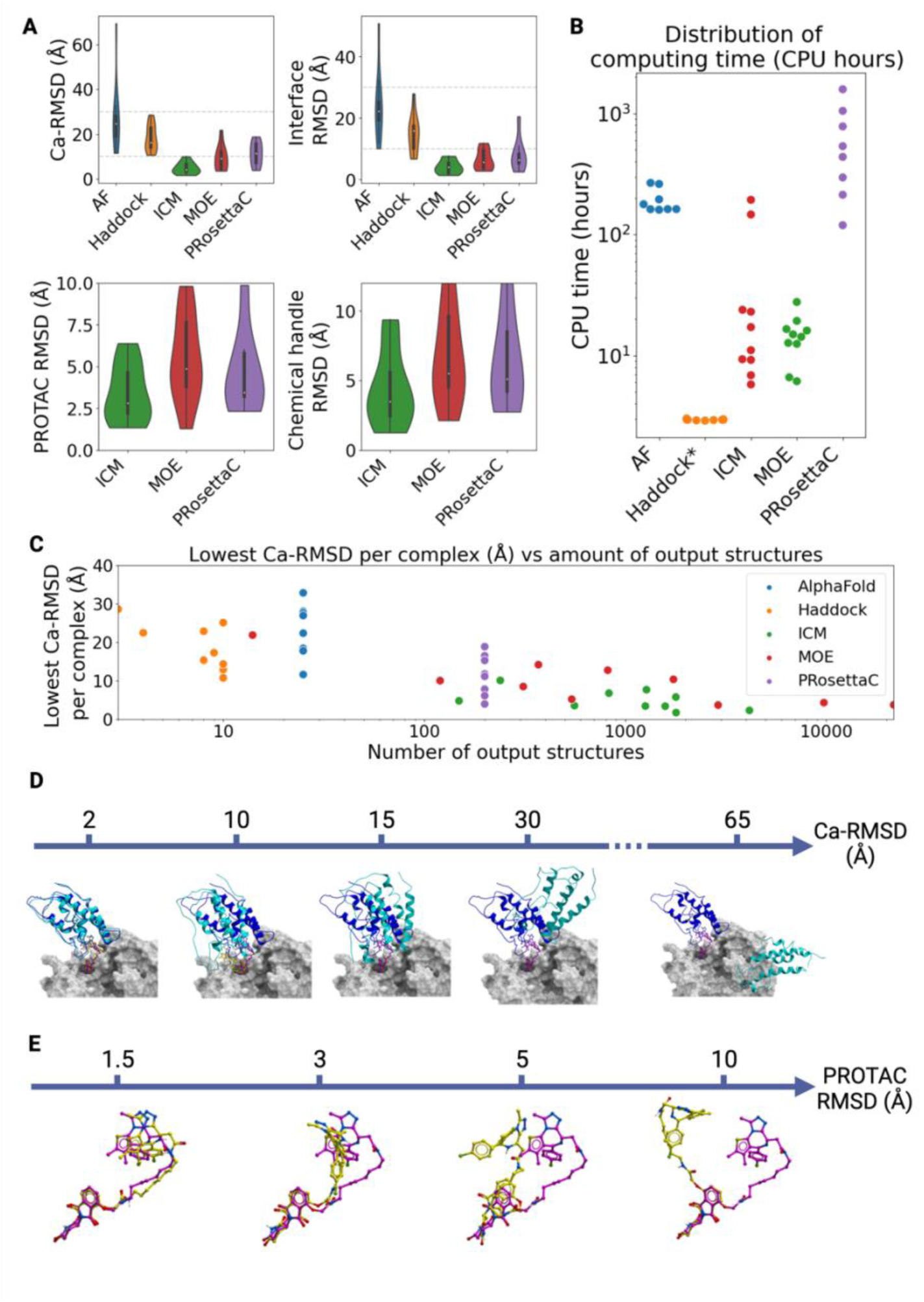
Structural accuracy of near-native ternary complex predictions. A) Distribution of the RMSD (Å) between the most accurate (near native) prediction and the crystal structure across the 10 complexes listed in Table 1, measured after superimposing the E3 ligase. Top-left: Ca-RMSD between the docked and crystallized target protein. Top-right: interface RMSD between the docked and crystallized target protein. Interface residues are defined as residues within <10Å of the E3 ligase. Bottom-left: RMSD between the docked and crystallized PROTAC. Bottom-right: RMSD between the docked and crystallized chemical handle of the target protein. B) Distribution of computing time per simulation displayed in log(CPU hours). C) Number of predicted structures for each simulation (i.e. each complex and method) plotted versus the lowest Ca-RMSD (Å) achieved by the simulation. D,E) Visual cue for RMSD values (Å). Protein and PROTAC crystal structures are shown in dark blue and pink respectively. Predictions are light blue and yellow. *Haddock is run on a webserver (log(simulation time)). **All methods use CPU except AlphaFold which requires a 1GPU:8CPU ratio. GPU hours not included.

**Table 3.** Average docking accuracy and compute time of the tested methods. Rank (%) is calculated as rank/(total number of predictions)*100. *AlphaFold requires 8 CPU for 1 GPU. GPU time is not incorporated. Other methods do not utilize GPUs. **Haddock is run on the webserver; time displayed is simulation hours and log(simulation hours).

| Method | Average Ca-RMSD (Å) of near-native complex | Average rank (%) of near-native complex | Average Ca-RMSD (Å) of top-ranked complex | Average Ca-RMSD (Å) of best in top 3 ranked complexes | Average number of output structures | Average computing time [CPU hours] |
| --- | --- | --- | --- | --- | --- | --- |
| AlphaFold | 28 Å | 48% | 39 Å | 36 Å | 25 | 193* |
| Haddock | 18 Å | 65% | 28 Å | 26 Å | 8 | 3** |
| MOE | 9 Å | 54% | 17 Å | 17 Å | 3637 | 45 |
| Molsoft | 5 Å | 29% | 26 Å | 25 Å | 1348 | 14 |
| PRosettaC | 11 Å | 57% | 29 Å | 30 Å | 200 | 431 |

The near-native complex is ranked on average in the top 29% of the ensemble of conformations predicted by ICM, which is better than other tools (> top 45%), but still far from perfect (Tab. 3). Ideally, a correlation should be observed where top-ranked structures have the lowest Ca-RMSD, but we rarely see such trend (Fig. 3). We note that the top-ranked conformation predicted with MOE is on average more accurate (17 Å Ca-RMSD) (Tab. 3), but it is questionable that this level of accuracy is sufficient for structure-based PROTAC design. Importantly, one may also ask whether the protein-protein interface captured crystallographically is the only conformational state stabilized by the PROTAC and is indeed a (or the only) conformation that productively induces the ubiquitination of the target protein (which we discuss later). It would be then conceivable that predicted ternary complex structures that significantly deviate from crystal structures are functionally relevant and may be used to predict active PROTACs. A corollary is that tools that perform poorly in replicating crystal structures may still be useful for PROTAC virtual screening.

**Figure 3.**
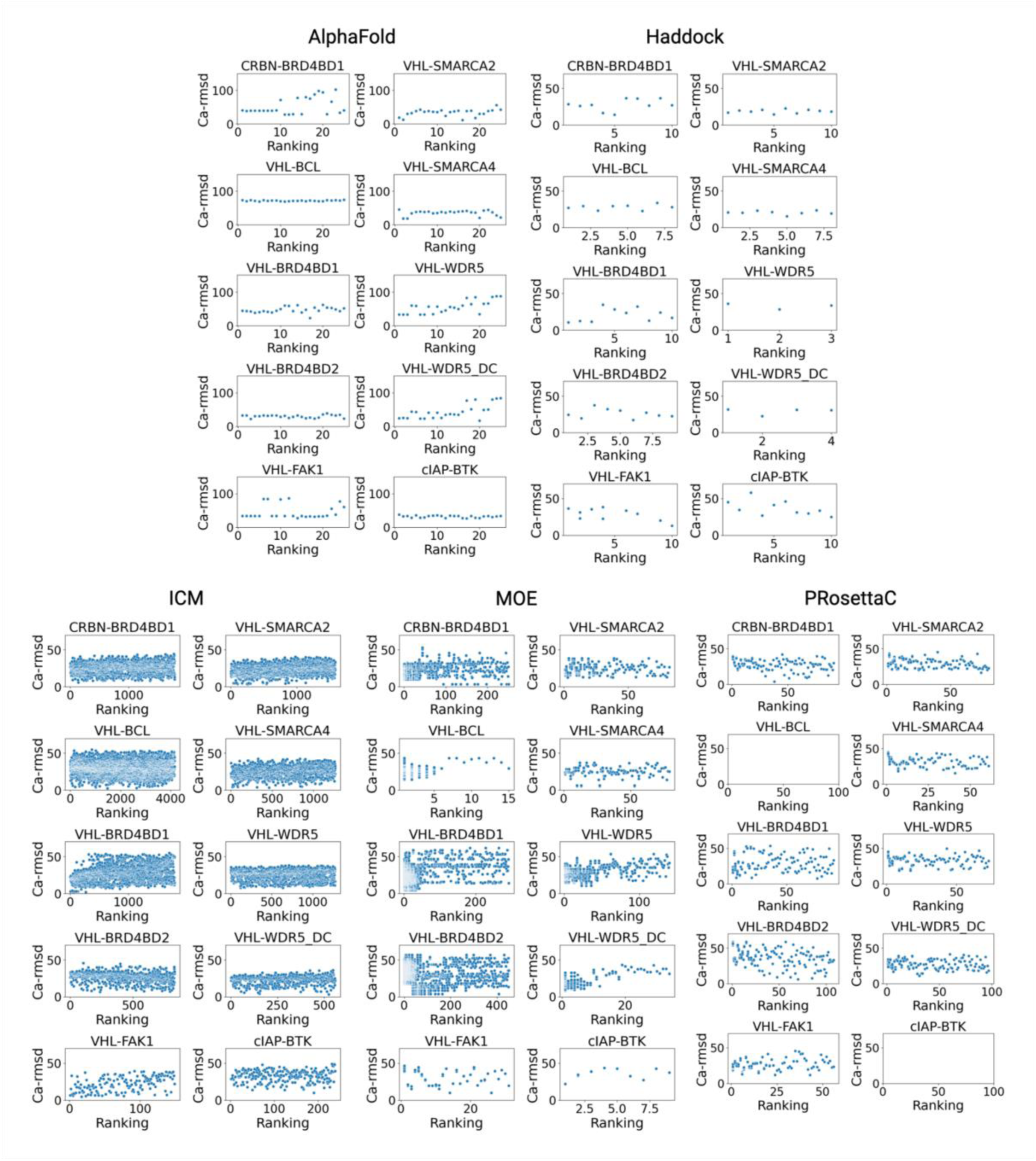
Correlation between rank of predicted structures and their Ca-RMSD with crystal structures. The ternary complex is superimposed on the E3 ligase and the Ca-RMSD is calculated for the target protein. For MOE and PRosettaC, different colors represent distinct conformational clusters.

### PROTAC virtual screening

#### Dissociating active from inactive PROTACs

The purpose of PROTAC virtual screening is to discriminate between active and inactive PROTACs which can be assessed by calculating the enrichment in active PROTACs among top-ranking molecules. We evaluated the selection of the top compound, top 3 or top 5 PROTACs (Fig. 4). Overall, the performance varies across test cases and methods. Encouragingly, the top-ranking molecule selected with MOE and ICM was active for four out of five test cases. MOE made a wrong prediction in what seemed the easiest challenge (8 out of 13 PARP1 PROTACs screened were active), while ICM failed the hardest challenge by selecting one of the 15 inactive CRBN-CDK2 PROTACs out 20 molecules screened. MOE performed slightly better than other tools when selecting three molecules, with around three-fold enrichment for three out of five test cases while enrichment values were less than two-fold with ICM and PRosettaC. Across methods and selection criteria, predictions were most accurate on the CRBN-CDK9 complex. On the other hand, all the predictions for the CRBN-PARP1 pair were poor regardless of method or selection. The exit vector of the crystallized chemical handle in the PARP1 structure is solvent-accessible but located deep in the binding pocket. A PROTAC linker must therefore first find its way out of the pocket, which is likely to be associated with severe conformational constraints. Finally, we found that in some cases conformational constraints on the chemical handles near the linker attachment point prevented proper superimposition of PROTACs to both handles. This led to dissociation of the CRBN-BRAF complex with ICM or aborted simulations of the CRBN-CDK complexes with PRosettaC. In both cases, reducing the number of atoms defining the chemical handle near the attachment point solved the issue.

**Figure 4.**
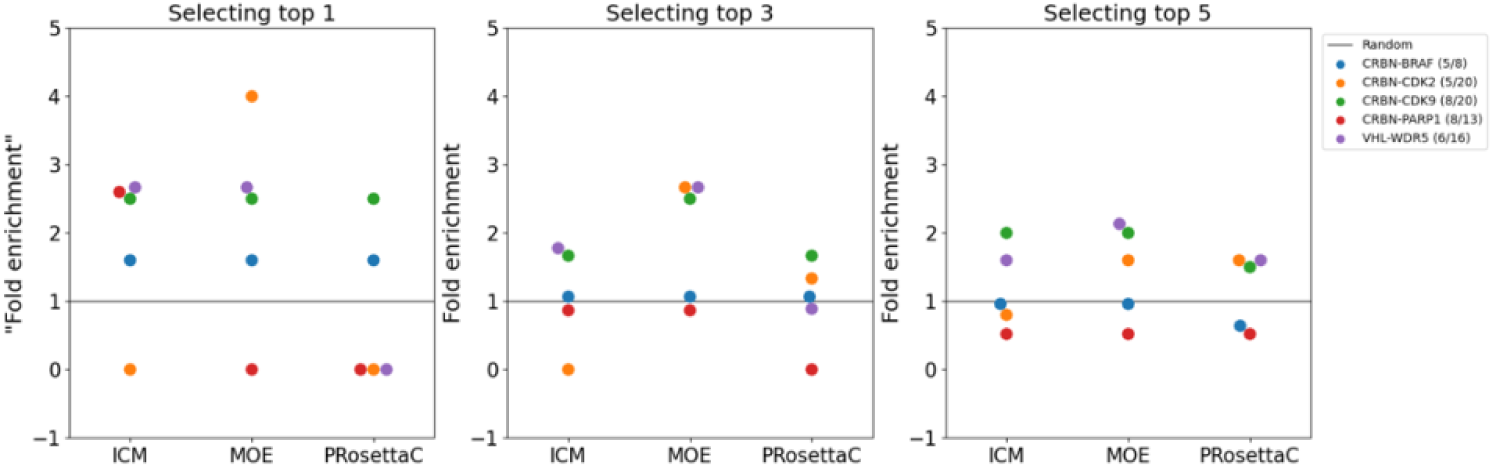
Fold enrichment when selecting the top, top 3 or top 5 predicted compound(s). Fold enrichment of 1 = random. The number of active PROTACs (active/total) are shown in the legend for each test case.

Considering the relatively low enrichment of active PROTACs among top-ranked compounds, we explored potential strategies to improve PROTAC virtual screening efficiency. We investigated whether defining putative ubiquitation zones on the target protein could serve as a beneficial filter, asked whether active PROTACs induced conformations not observed with inactive PROTACs and used the crystal structure of a ternary complex as input for PROTAC virtual screening.

### New metrics to improve PROTAC virtual screening efficiency

#### Searching for optimal ubiquitination zones

The E3-PROTAC-Target complex is part of a multiprotein complex responsible for the transfer of ubiquitin to the target protein. In Cullin RING E3 ligases, the complex consists of the E3 ligase, a one of four Cullin proteins, adaptor proteins and the E2 ligase conjugated to ubiquitin (Fig. 5). The acceptable distance between the E2 ligase and the substrate lysine, which defines what we call here the ubiquitination zone, may be critical for successful ubiquitin transfer and may be used as a metric to filter-out non-productive ternary complex structures. To determine the ubiquitination zone, we superimposed multiple crystal structures of a given ligase in complex with different target proteins and measured the distance between lysines known to be ubiquitinated according to the PhosphoSitePlus database and the E2 ligase-binding adapter protein RBX1. The E2 ligase is not crystallized in the complex, so the distance to its adapter protein is used instead. Only crystal structures with full-length target proteins were included, because structures of isolated domains do not include all the lysines residues known to be ubiquitinated for a protein. Since CRBN E3 ligase is only crystallized with PROTACs that target isolated domains, the analysis could only be performed with the VHL E3 ligase. After superimposing WDR5 (PDB: 7JTO^21^, 7JTP^21^, 7Q2J^30^, 8BB2, 8BB3, 8BB4, 8BB5) and Bcl-xL (PDB: 6ZHC^16^) on the VHL-Cullin-2-RBX1 complex, all potential lysine substrates resided within 30Å to 80Å of RBX1 (Fig. 5). We hypothesized that, while it may vary from one E3 complex to another, the ubiquitination zone for a given E3 would be a smaller range within this extended region of 30Å-80Å, and may be predictive of PROTAC activity. We superimposed all the ternary complexes generated during the PROTAC virtual screening (ICM) onto the composite cullin RING crystal structure and determined the number of lysine residues in a series of ubiquitination zones each spanning 20Å. Both the percentage of structures with a lysine residue (listed in PhosphoSitePlus) in the ubiquitination zone and the number of lysines available in the ubiquitination zone were determined for each PROTAC. We observed no correlation between either metric and PROTAC activity (Fig. S2-3). The lack of correlation could be attributed to the documented flexibility of the U-shaped cullin-RING complexes which allows for ubiquitination of substrates in wide spatial range.

**Figure 5.**
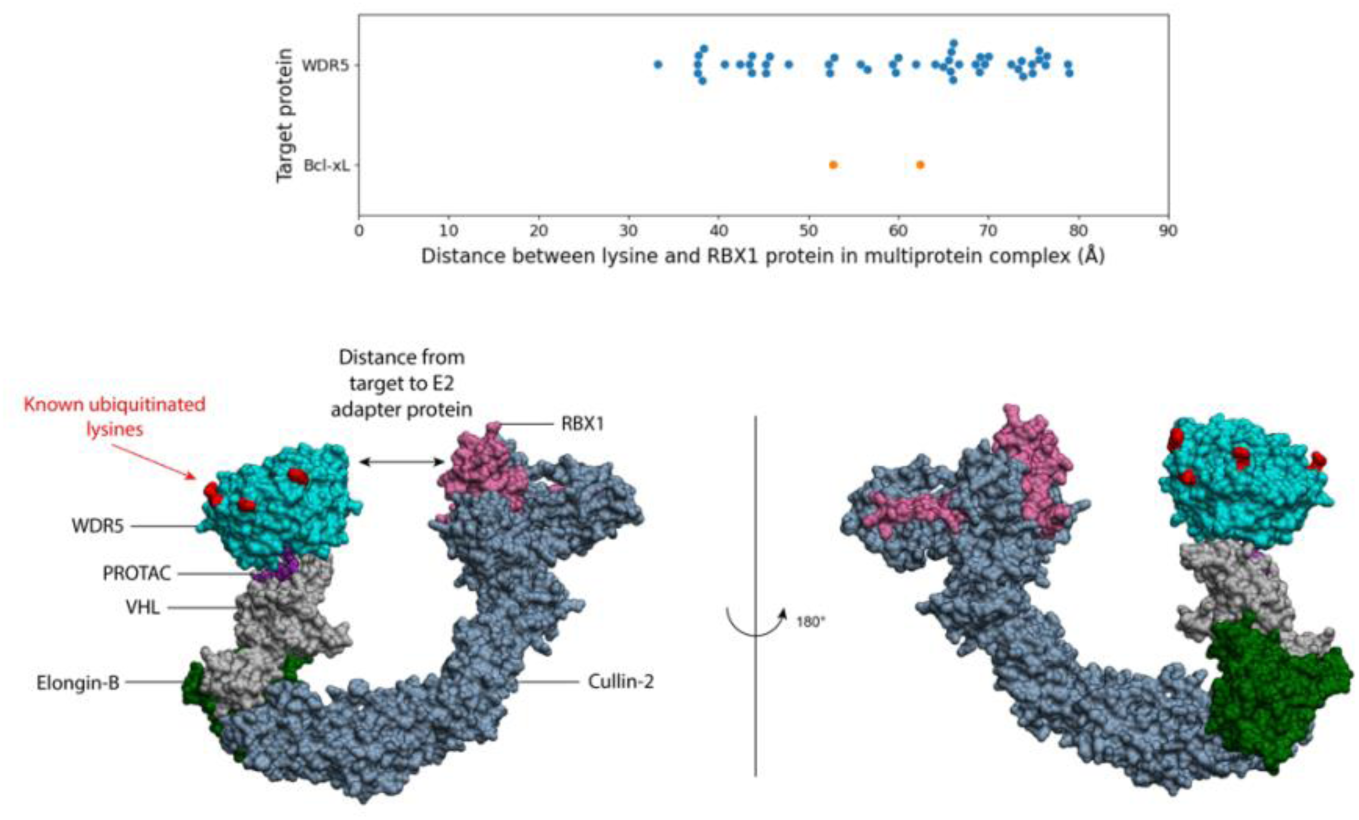
Minimum distance between ubiquitinated or acetylated lysine residues on the target proteins and RBX1 protein in the multiprotein complex. Composite structures of the RBX1-CUL2-VHL-WDR5 complex (bottom panel, PDB codes 5N4W^29^ and 7JTP^21^ (other complexes not shown, PDB codes 7JTO^21^, 7Q2J^30^, 8BB2, 8BB3, 8BB4, 8BB5)), and the RBX1-CUL2-VHL-Bcl-xL complex (not shown, PDB codes 5N4W^29^ and 6ZHC^16^) were used to measure distances separating ubiquitinated lysines and RBX1 (top panel).

#### Looking for conformations that dissociate active from inactive PROTACs

Active PROTACs are expected to induce ternary complex conformations that inactive PROTACs cannot induce. We looked for conformational clusters that were highly enriched in active PROTACs and devoid of inactive PROTACs that may be used to prioritize follow-up molecules. Encouragingly, we found for each test case an “actives-only” cluster (a ternary complex conformation only generated with active PROTACs). For the CRBN-BRAF complex, all active PROTACs could be found in actives-only clusters. In fact, one cluster was induced by three of the five active PROTACs and none of the 3 inactives (Fig. 6). In other test cases, 40% to 83% of actives shared a ternary complex conformation absent of inactive molecules (Tab. S9). However, due to limitations in available data, the statistical significance of these observations remains weak. Indeed, in all test cases, no more than three PROTACs are found in any given cluster. The restriction of this approach is that prior data on active and inactive PROTACs is required to find actives-only conformational clusters. We note that the conformational arrangements in the actives-only VHL-WDR5 clusters differed from the crystal structures of the complex (PDB: 7JTP^21^ and 7Q2J^30^), with Ca-RMSD >10 Å in all cases (Fig. 6), which could either raise a red flag on this approach, or support the notion that crystal structures do not necessarily reflect the only possible conformational arrangement.

**Figure 6.**
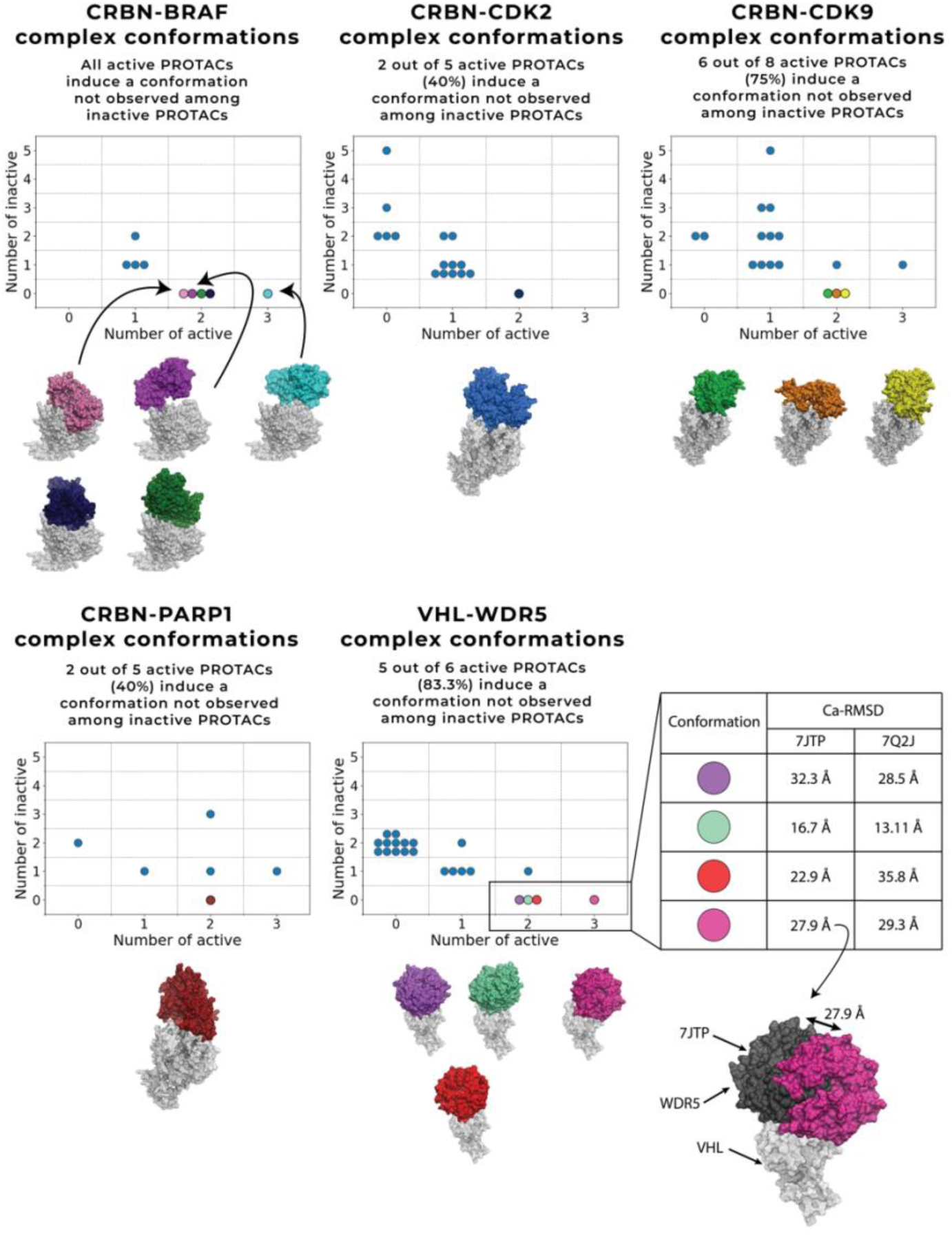
Can some ternary complex conformations dissociate active from inactive PROTACs. Each dot represents a protein-protein complex conformational cluster. Conformations only observed among active PROTACs are highlighted with different colors. Ca-RMSD between active-only clusters and crystal structures are provided for the VHL-WDR5 complex.

#### Virtual screening using the crystal structure

Gadd et al. reported that the crystal structure of a ternary complex with an active PROTAC could be used to design optimized molecules^13^. To evaluate whether virtual screening against the crystal structures of the protein-protein interface could more efficiently dissociate active from inactive PROTACs, we selected an option available in MOE to use the crystallized forms of the VHL-WDR5 complex (PDB: 7JTP^21^ and 7Q2J^30^) instead of computationally docked protein-protein interfaces, as input for virtual screening. The other test cases with available screening data did not have a crystal structure of the ternary complex. Strikingly, results were worse when using crystal structures than using docked proteins: no active PROTAC was found in the top 3 ranked compounds and only one in the top 5 (Fig. 7), while the top-ranked molecule selected with the docked protein-protein interface was active (Fig. 6). It is dangerous to draw conclusions based on a single case study, but this result does not support the notion that using a crystal structure increases the efficiency of PROTAC virtual screening.

**Figure 7.**
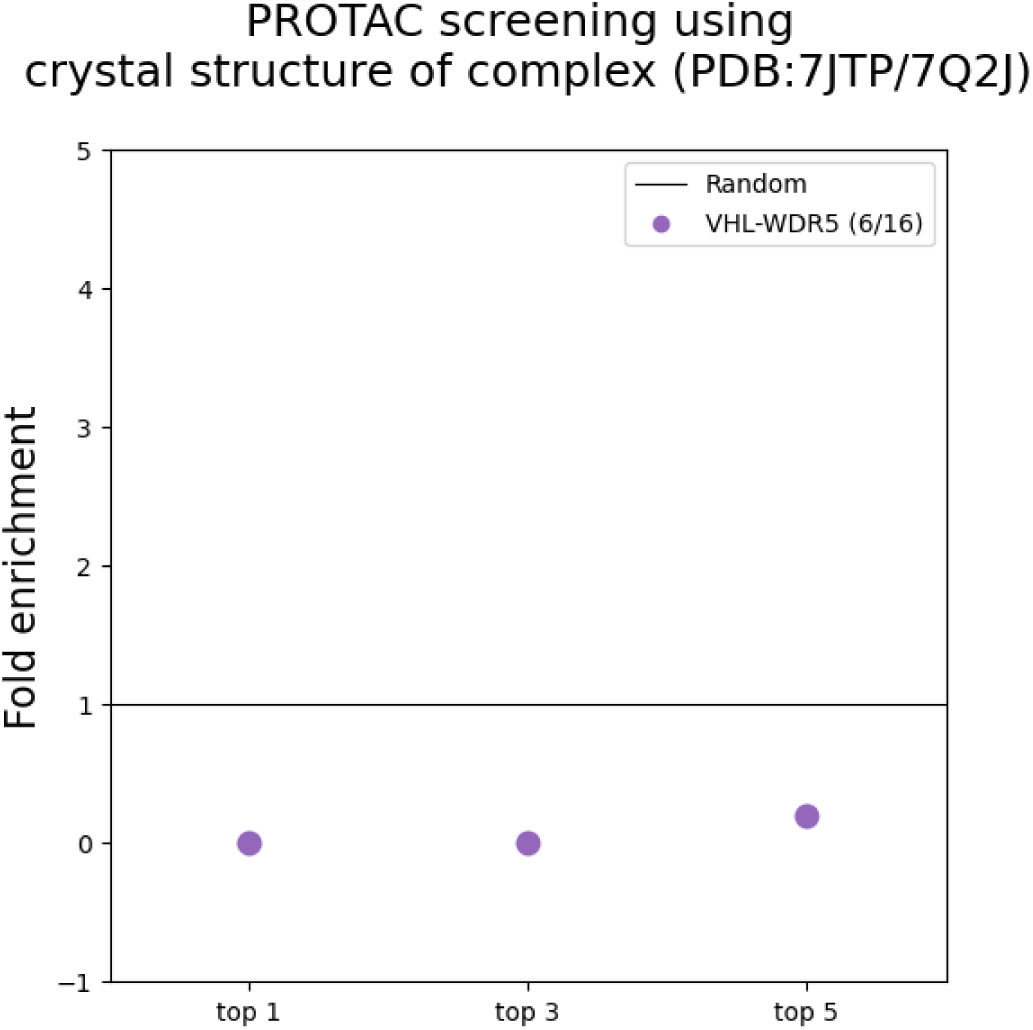
Enrichment in active PROTACs using crystal structures of the ternary complex as input. Virtual screening using the crystal structures of the VHL-WDR5 complex as input in the MOE protocol (PDB: 7JTP^21^/7Q2J^30^). Fold enrichment of 1 = random. The number of active compounds (active/total) is provided in the legend.

## Discussion

Targeted protein degradation is a promising modality for cancer, immune and neurodegenerative diseases and numerous PROTACs are now in clinical trials^1^. The discovery and optimization of PROTACs is mostly a trial-and-error process guided more by medicinal chemistry intuition than structure-based design strategies. Here, we systematically benchmarked three computational methods specifically customized to predict the structure of ternary complexes and virtually screen PROTAC libraries: ICM, MOE and PRosettaC. This field is evolving rapidly, and new screening methods are continuously emerging. Each method uses different principles and techniques to create ternary complexes or virtually screen PROTAC candidates.

In this benchmark, protein structures in their unbound state (structures without bound PROTAC or protein partner) are used. We generally observe better predictions (i.e. complex conformations closer to the crystal structures) using structures from the crystallized complex as input. This is to be expected as sidechains and sometimes even entire loops adopt different conformations in the bound and unbound state, as seen in the VHL-SMARCA2 complex^19^. While the computational methods account for side-chain flexibility, the protein backbone is kept rigid. Future improvements in screening methods should therefore better take into consideration the dynamic nature of the complex.

We show that PROTAC virtual screening does not improve when using the crystal structure of the ternary complex. This questions whether the crystal structure is the only and relevant ternary complex. Observations that support this idea are that 1) conformations induced only by active PROTACs are not similar to crystal structures, 2) near-native models rank badly during the simulations, 3) different ternary complex conformations are observed with different WDR5 PROTACs (PDB: 7JTP^21^, 7JTO^21^ form a different complex than PDB: 7Q2J^30^, 8BB2, 8BB3, 8BB4, 8BB5), and lastly 4) other studies support the idea of multiple conformations of the ternary complex. The study by Villegas et al. shows that multiple structural arrangements are associated with local energy minim. It also highlights that the crystal structures of the ternary complexes is not always the most favourable complex in terms of energy or solvation potential^32^. The study by Schwalm et al. showed that inactive PROTACs still form ternary complexes *in vitro*.^33^ This supports the notion that active PROTACs induce a specific ternary complex that results in ubiquitination which could not be accessed by inactive PROTACs. Overall, the crystal structures are a snapshot of a dynamic complex, therefore more dynamic approaches like molecular dynamics or cryo-em could potentially explore the conformational space of PROTAC ternary complex, as recently conducted by Liao et al.^34^.

While we were encouraged to see that the top ranked compound selected by MOE and ICM was almost systematically active, results were less clear for the top 3 or top 5 ranked compounds, and critically, the statistical relevance of these results was weak due to the limited size of the dataset. Clearly, more experimental data on active and especially inactive PROTACs is needed to develop and evaluate PROTAC screening methods.

We tested different protocols to improve PROTAC screening efficiency. We find that filtering out PROTACs that fail to place substrate lysines inside a ubiquitination zone delimited by a specific distance from the E2 protein does not improve predictions. We note that we ignored the potential dynamics of this large multiprotein degradation complex in this exercise while MD simulations showed that the probability density of a lysine in a ubiquitination zone is indicative of activity^26^. More encouragingly, we found that ternary complex conformations could often be found that were exclusively composed of active PROTACS. However, due to limited data, these results are not significant and need to be explored further.

## Conclusion

Computational prediction of PROTAC ternary complex structures and PROTAC virtual screening may be seen as a daunting task. Pioneer efforts to develop tools that meet this challenge are commendable. By benchmarking some of these tools, we hope to have delineated limitations, reasonable expectations, and possible path for improvement. Without a significant increase in available experimental data, improvements in the field will be slow and limited: we urge experimentalists to better share their structure-activity relationships, including inactive PROTACs.

## Data availability

The data associated with this study are available at a Zenodo (https://zenodo.org/record/8298749) repository. The repository includes all the input files. Researchers can access and download the data to reproduce the results presented in this manuscript. The raw data files are available on request. The software is available at ICM (https://molsoft.com/), MOE (https://www.chemcomp.com/), PRosettaC (https://github.com/LondonLab/PRosettaC / https://hub.docker.com/r/erovers/prosettac), Haddock (https://wenmr.science.uu.nl/haddock2.4/) and AlphaFold (https://github.com/deepmind/alphafold).

## Acknowledgements

This research was enabled in part by support provided by Compute Ontario (https://www.computeontario.ca/) and the Digital Research Alliance of Canada (alliancecan.ca). Figures were created using Biorender (www.BioRender.com). MS gratefully acknowledges support from NSERC [Grant RGPIN-2019-04416], CQDM (Quantum Leap-176) and MITACS accelerate (IT13051). The Structural Genomics Consortium is a registered charity (no: 1097737) that receives funds from Bayer AG, Boehringer Ingelheim, Bristol Myers Squibb, Genentech, Genome Canada through Ontario Genomics Institute [OGI-196], EU/EFPIA/OICR/McGill/KTH/Diamond Innovative Medicines Initiative 2 Joint Undertaking [EUbOPEN grant 875510], Janssen, Merck KGaA (aka EMD in Canada and US), Pfizer and Takeda.

## References

1. Békés, M.; Langley, D. R.; Crews, C. M. PROTAC Targeted Protein Degraders: The Past Is Prologue. Nat. Rev. Drug Discov. 2022, 21, 181–200. 10.1038/s41573-021-00371-6.

2. Modell, A. E.; Lai, S.; Nguyen, T. M.; Choudhary, A. Bifunctional Modalities for Repurposing Protein Function. Cell Chem. Biol. 2021, 28 (7), 1081–1089. 10.1016/j.chembiol.2021.06.005.

3. Hanzl, A.; Winter, G. E. Targeted Protein Degradation: Current and Future Challenges. Curr. Opin. Chem. Biol. 2020, 56, 35–41. 10.1016/j.cbpa.2019.11.012.

4. He, Y.; Zhang, X.; Chang, J.; Kim, H.-N.; Zhang, P.; Wang, Y.; Khan, S.; Liu, X.; Zhang, X.; Lv, D.; Song, L.; Li, W.; Thummuri, D.; Yuan, Y.; Wiegand, J. S.; Ortiz, Y. T.; Budamagunta, V.; Elisseeff, J. H.; Campisi, J.; Almeida, M.; Zheng, G.; Zhou, D. Using Proteolysis-Targeting Chimera Technology to Reduce Navitoclax Platelet Toxicity and Improve Its Senolytic Activity. Nat. Commun. 2020, 11 (1), 1996. 10.1038/s41467-020-15838-0.

5. Hendrick, C. E.; Jorgensen, J. R.; Chaudhry, C.; Strambeanu, I. I.; Brazeau, J. F.; Schiffer, J.; Shi, Z.; Venable, J. D.; Wolkenberg, S. E. Direct-to-Biology Accelerates PROTAC Synthesis and the Evaluation of Linker Effects on Permeability and Degradation. ACS Med. Chem. Lett. 2022, 13 (7), 1182–1190. 10.1021/ACSMEDCHEMLETT.2C00124.

6. Tran, N. L.; Leconte, G. A.; Ferguson, F. M. Targeted Protein Degradation: Design Considerations for PROTAC Development. Curr. Protoc. 2022, 2 (12). 10.1002/CPZ1.611.

7. Zaidman, D.; Prilusky, J.; London, N. PRosettaC: Rosetta Based Modeling of PROTAC Mediated Ternary Complexes. J. Chem. Inf. Model. 2020, 60 (10), 4894–4903. 10.1021/acs.jcim.0c00589.

8. Drummond, M. L.; Williams, C. I. In Silico Modeling of PROTAC-Mediated Ternary Complexes: Validation and Application. J. Chem. Inf. Model. 2019, 59 (4), 1634–1644. 10.1021/acs.jcim.8b00872.

9. Drummond, M. L.; Henry, A.; Li, H.; Williams, C. I. Improved Accuracy for Modeling PROTAC-Mediated Ternary Complex Formation and Targeted Protein Degradation via New In Silico Methodologies. J. Chem. Inf. Model. 2020, 60 (10), 5234–5254. 10.1021/acs.jcim.0c00897.

10. Dominguez, C.; Boelens, R.; Bonvin, A. M. J. J. HADDOCK: A Protein−Protein Docking Approach Based on Biochemical or Biophysical Information. J. Am. Chem. Soc. 2003, 125 (7), 1731–1737. 10.1021/ja026939x.

11. Evans, R.; O’Neill, M.; Pritzel, A.; Antropova, N.; Senior, A.; Green, T.; Žídek, A.; Bates, R.; Blackwell, S.; Yim, J.; Ronneberger, O.; Bodenstein, S.; Zielinski, M.; Bridgland, A.; Potapenko, A.; Cowie, A.; Tunyasuvunakool, K.; Jain, R.; Clancy, E.; Kohli, P.; Jumper, J.; Hassabis, D. Protein Complex Prediction with AlphaFold-Multimer. bioRxiv 2022, 2021.10.04.463034. 10.1101/2021.10.04.463034.

12. Nowak, R. P.; Deangelo, S. L.; Buckley, D.; He, Z.; Donovan, K. A.; An, J.; Safaee, N.; Jedrychowski, M. P.; Ponthier, C. M.; Ishoey, M.; Zhang, T.; Mancias, J. D.; Gray, N. S.; Bradner, J. E.; Fischer, E. S. Plasticity in Binding Confers Selectivity in Ligand-Induced Protein Degradation. Nat. Chem. Biol. 2018, 14 (7), 706–714. 10.1038/S41589-018-0055-Y.

13. Gadd, M. S.; Testa, A.; Lucas, X.; Chan, K. H.; Chen, W.; Lamont, D. J.; Zengerle, M.; Ciulli, A. Structural Basis of PROTAC Cooperative Recognition for Selective Protein Degradation. Nat. Chem. Biol. 2017 135 2017, 13 (5), 514–521. 10.1038/nchembio.2329.

14. Rovers, E.; Schapira, M. Methods for Computer-Assisted PROTAC Design; Methods in Enzymology; Academic Press, 2023. 10.1016/bs.mie.2023.06.020.

15. Schiemer, J.; Horst, R.; Meng, Y.; Montgomery, J. I.; Xu, Y.; Feng, X.; Borzilleri, K.; Uccello, D. P.; Leverett, C.; Brown, S.; Che, Y.; Brown, M. F.; Hayward, M. M.; Gilbert, A. M.; Noe, M. C.; Calabrese, M. F. Snapshots and Ensembles of BTK and CIAP1 Protein Degrader Ternary Complexes. Nat. Chem. Biol. 2021, 17 (2), 152–160. 10.1038/S41589-020-00686-2.

16. Chung, C. W.; Dai, H.; Fernandez, E.; Tinworth, C. P.; Churcher, I.; Cryan, J.; Denyer, J.; Harling, J. D.; Konopacka, A.; Queisser, M. A.; Tame, C. J.; Watt, G.; Jiang, F.; Qian, D.; Benowitz, A. B. Structural Insights into PROTAC-Mediated Degradation of Bcl-XL. ACS Chem. Biol. 2020, 15 (9), 2316–2323. 10.1021/ACSCHEMBIO.0C00266.

17. Dragovich, P. S.; Pillow, T. H.; Blake, R. A.; Sadowsky, J. D.; Adaligil, E.; Adhikari, P.; Chen, J.; Corr, N.; Dela Cruz-Chuh, J.; Del Rosario, G.; Fullerton, A.; Hartman, S. J.; Jiang, F.; Kaufman, S.; Kleinheinz, T.; Kozak, K. R.; Liu, L.; Lu, Y.; Mulvihill, M. M.; Murray, J. M.; O’Donohue, A.; Rowntree, R. K.; Sawyer, W. S.; Staben, L. R.; Wai, J.; Wang, J.; Wei, B.; Wei, W.; Xu, Z.; Yao, H.; Yu, S. F.; Zhang, D.; Zhang, H.; Zhang, S.; Zhao, Y.; Zhou, H.; Zhu, X. Antibody-Mediated Delivery of Chimeric BRD4 Degraders. Part 2: Improvement of In Vitro Antiproliferation Activity and In Vivo Antitumor Efficacy. J. Med. Chem. 2021, 64 (5), 2576–2607. 10.1021/ACS.JMEDCHEM.0C01846.

18. Law, R. P.; Nunes, J.; Chung, C. wa; Bantscheff, M.; Buda, K.; Dai, H.; Evans, J. P.; Flinders, A.; Klimaszewska, D.; Lewis, A. J.; Muelbaier, M.; Scott-Stevens, P.; Stacey, P.; Tame, C. J.; Watt, G. F.; Zinn, N.; Queisser, M. A.; Harling, J. D.; Benowitz, A. B. Discovery and Characterisation of Highly Cooperative FAK-Degrading PROTACs. Angew. Chem. Int. Ed. Engl. 2021, 60 (43), 23327–23334. 10.1002/ANIE.202109237.

19. Farnaby, W.; Koegl, M.; Roy, M. J.; Whitworth, C.; Diers, E.; Trainor, N.; Zollman, D.; Steurer, S.; Karolyi-Oezguer, J.; Riedmueller, C.; Gmaschitz, T.; Wachter, J.; Dank, C.; Galant, M.; Sharps, B.; Rumpel, K.; Traxler, E.; Gerstberger, T.; Schnitzer, R.; Petermann, O.; Greb, P.; Weinstabl, H.; Bader, G.; Zoephel, A.; Weiss-Puxbaum, A.; Ehrenhöfer-Wölfer, K.; Wöhrle, S.; Boehmelt, G.; Rinnenthal, J.; Arnhof, H.; Wiechens, N.; Wu, M.-Y.; Owen-Hughes, T.; Ettmayer, P.; Pearson, M.; McConnell, D. B.; Ciulli, A. BAF Complex Vulnerabilities in Cancer Demonstrated via Structure-Based PROTAC Design. Nat. Chem. Biol. 2019, 15 (7), 672–680. 10.1038/s41589-019-0294-6.

20. Schwalm, M. P.; Krämer, A.; Dölle, A.; Weckesser, J.; Yu, X.; Jin, J.; Saxena, K.; Knapp, S. Tracking the PROTAC Degradation Pathway in Living Cells Highlights the Importance of Ternary Complex Measurement for PROTAC Optimization. bioRxiv 2023, 2023.01.11.523589. 10.1101/2023.01.11.523589.

21. Yu, X.; Li, D.; Kottur, J.; Shen, Y.; Kim, H. S.; Park, K. S.; Tsai, Y. H.; Gong, W.; Wang, J.; Suzuki, K.; Parker, J.; Herring, L.; Kaniskan, H. Ü.; Cai, L.; Jain, R.; Liu, J.; Aggarwal, A. K.; Wang, G. G.; Jin, J. A Selective WDR5 Degrader Inhibits Acute Myeloid Leukemia in Patient-Derived Mouse Models. Sci. Transl. Med. 2021, 13 (613). 10.1126/SCITRANSLMED.ABJ1578.

22. Posternak, G.; Tang, X.; Maisonneuve, P.; Jin, T.; Lavoie, H.; Daou, S.; Orlicky, S.; Goullet de Rugy, T.; Caldwell, L.; Chan, K.; Aman, A.; Prakesch, M.; Poda, G.; Mader, P.; Wong, C.; Maier, S.; Kitaygorodsky, J.; Larsen, B.; Colwill, K.; Yin, Z.; Ceccarelli, D. F.; Batey, R. A.; Taipale, M.; Kurinov, I.; Uehling, D.; Wrana, J.; Durocher, D.; Gingras, A. C.; Al-Awar, R.; Therrien, M.; Sicheri, F. Functional Characterization of a PROTAC Directed against BRAF Mutant V600E. Nat. Chem. Biol. 2020, 16 (11), 1170–1178. 10.1038/S41589-020-0609-7.

23. Zhou, F.; Chen, L.; Cao, C.; Yu, J.; Luo, X.; Zhou, P.; Zhao, L.; Du, W.; Cheng, J.; Xie, Y.; Chen, Y. Development of Selective Mono or Dual PROTAC Degrader Probe of CDK Isoforms. Eur. J. Med. Chem. 2020, 187. 10.1016/j.ejmech.2019.111952.

24. Wang, S.; Han, L.; Han, J.; Li, P.; Ding, Q.; Zhang, Q. J.; Liu, Z. P.; Chen, C.; Yu, Y. Uncoupling of PARP1 Trapping and Inhibition Using Selective PARP1 Degradation. Nat. Chem. Biol. 2019, 15 (12), 1223–1231. 10.1038/s41589-019-0379-2.

25. Dölle, A.; Adhikari, B.; Krämer, A.; Weckesser, J.; Berner, N.; Berger, L. M.; Diebold, M.; Szewczyk, M. M.; Barsyte-Lovejoy, D.; Arrowsmith, C. H.; Gebel, J.; Löhr, F.; Dötsch, V.; Eilers, M.; Heinzlmeir, S.; Kuster, B.; Sotriffer, C.; Wolf, E.; Knapp, S. Design, Synthesis, and Evaluation of WD-Repeat-Containing Protein 5 (WDR5) Degraders. J. Med. Chem. 2021, 64 (15), 10682–10710. 10.1021/acs.jmedchem.1c00146.

26. Dixon, T.; MacPherson, D.; Mostofian, B.; Dauzhenka, T.; Lotz, S.; McGee, D.; Shechter, S.; Shrestha, U. R.; Wiewiora, R.; McDargh, Z. A.; Pei, F.; Pal, R.; Ribeiro, J. V.; Wilkerson, T.; Sachdeva, V.; Gao, N.; Jain, S.; Sparks, S.; Li, Y.; Vinitsky, A.; Zhang, X.; Razavi, A. M.; Kolossváry, I.; Imbriglio, J.; Evdokimov, A.; Bergeron, L.; Zhou, W.; Adhikari, J.; Ruprecht, B.; Dickson, A.; Xu, H.; Sherman, W.; Izaguirre, J. A. Predicting the Structural Basis of Targeted Protein Degradation by Integrating Molecular Dynamics Simulations with Structural Mass Spectrometry. Nat. Commun. 2022 131 2022, 13 (1), 1–24. 10.1038/s41467-022-33575-4.

27. Hornbeck, P. V.; Zhang, B.; Murray, B.; Kornhauser, J. M.; Latham, V.; Skrzypek, E. PhosphoSitePlus, 2014: Mutations, PTMs and Recalibrations. Nucleic Acids Res. 2015, 43 (Database issue), D512–D520. 10.1093/NAR/GKU1267.

28. Zhang, W.; Roy Burman, S. S.; Chen, J.; Donovan, K. A.; Cao, Y.; Shu, C.; Zhang, B.; Zeng, Z.; Gu, S.; Zhang, Y.; Li, D.; Fischer, E. S.; Tokheim, C.; Shirley Liu, X. Machine Learning Modeling of Protein-Intrinsic Features Predicts Tractability of Targeted Protein Degradation. Genomics. Proteomics Bioinformatics 2022, 20 (5), 882–898. 10.1016/J.GPB.2022.11.008.

29. Cardote, T. A. F.; Gadd, M. S.; Ciulli, A. Crystal Structure of the Cul2-Rbx1-EloBC-VHL Ubiquitin Ligase Complex. Structure 2017, 25 (6), 901–911.e3. 10.1016/J.STR.2017.04.009.

30. Chen, P.-H.; Hu, Z.; An, E.; Okeke, I.; Zheng, S.; Luo, X.; Gong, A.; Jaime-Figueroa, S.; Crews, C. M. Modulation of Phosphoprotein Activity by Phosphorylation Targeting Chimeras (PhosTACs). ACS Chem. Biol. 2021. 10.1021/acschembio.1c00693.

31. Méndez, R.; Leplae, R.; Lensink, M. F.; Wodak, S. J. Assessment of CAPRI Predictions in Rounds 3–5 Shows Progress in Docking Procedures. Proteins Struct. Funct. Bioinforma. 2005, 60 (2), 150–169. 10.1002/PROT.20551.

32. Villegas, J. A.; Vaid, T. M.; Johnson, M. E.; Moore, T. W. Mapping the Energy Landscape of PROTAC-Mediated Protein-Protein Interactions. Comput. Struct. Biotechnol. J. 2023, 21, 1885–1892. 10.1016/J.CSBJ.2023.02.049.

33. Schwalm, M. P.; Krämer, A.; Dölle, A.; Weckesser, J.; Yu, X.; Jin, J.; Saxena, K.; Knapp, S. Tracking the PROTAC Degradation Pathway in Living Cells Highlights the Importance of Ternary Complex Measurement for PROTAC Optimization. Cell Chem. Biol. 2023, 753–765. 10.1016/j.chembiol.2023.06.002.

34. Liao, J.; Nie, X.; Unarta, I. C.; Ericksen, S. S.; Tang, W. In Silico Modeling and Scoring of PROTAC-Mediated Ternary Complex Poses. J. Med. Chem. 2022. 10.1021/acs.jmedchem.1c02155.

