## Supporting information for "Benchmarking of PROTAC docking and virtual screening tools"

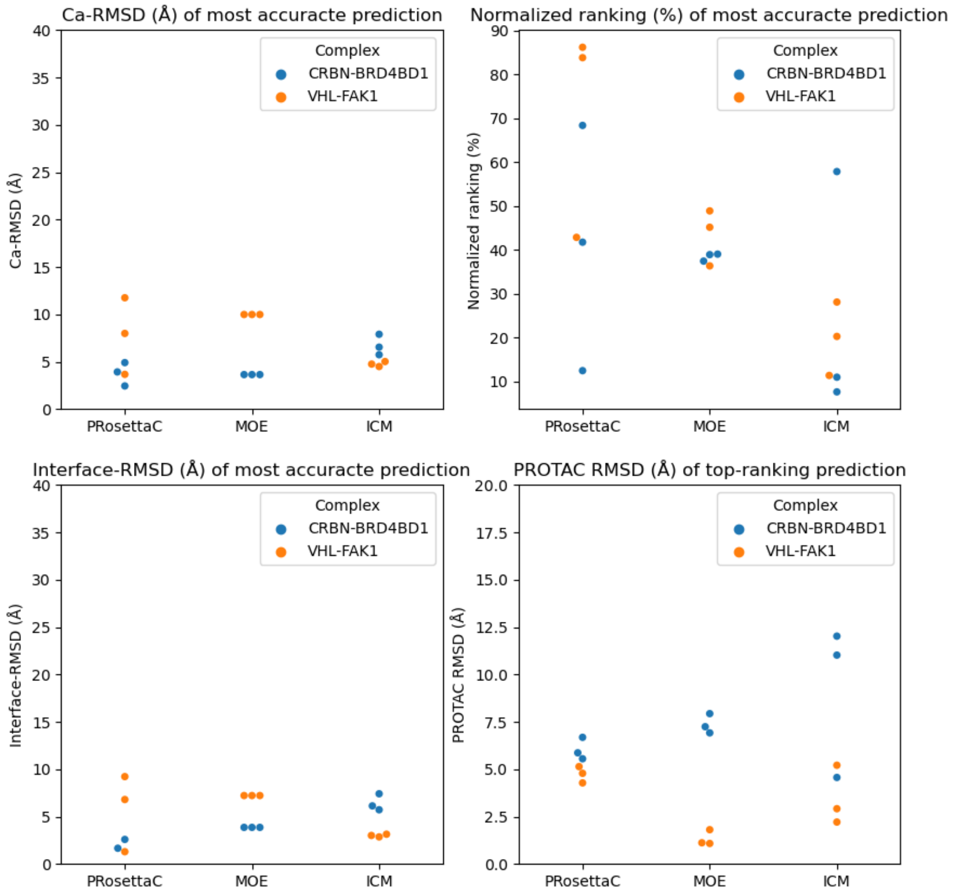

**Figure S1. Reproducibility of the docking simulations.** The simulations are repeated in triplicate with each tool for CRBN-BRD4BD1 (A) and VHL-FAK1 (B) and results are compared with the crystal structure. In each case, the Ca-RMSD (top-left) and rank (top-right) of the most accurate (near-native) conformation are determined, as well as the Ca-RMSD of the top-ranking conformation (bottom). Ranking is normalized by rank/(total number of predicted conformations)*100.

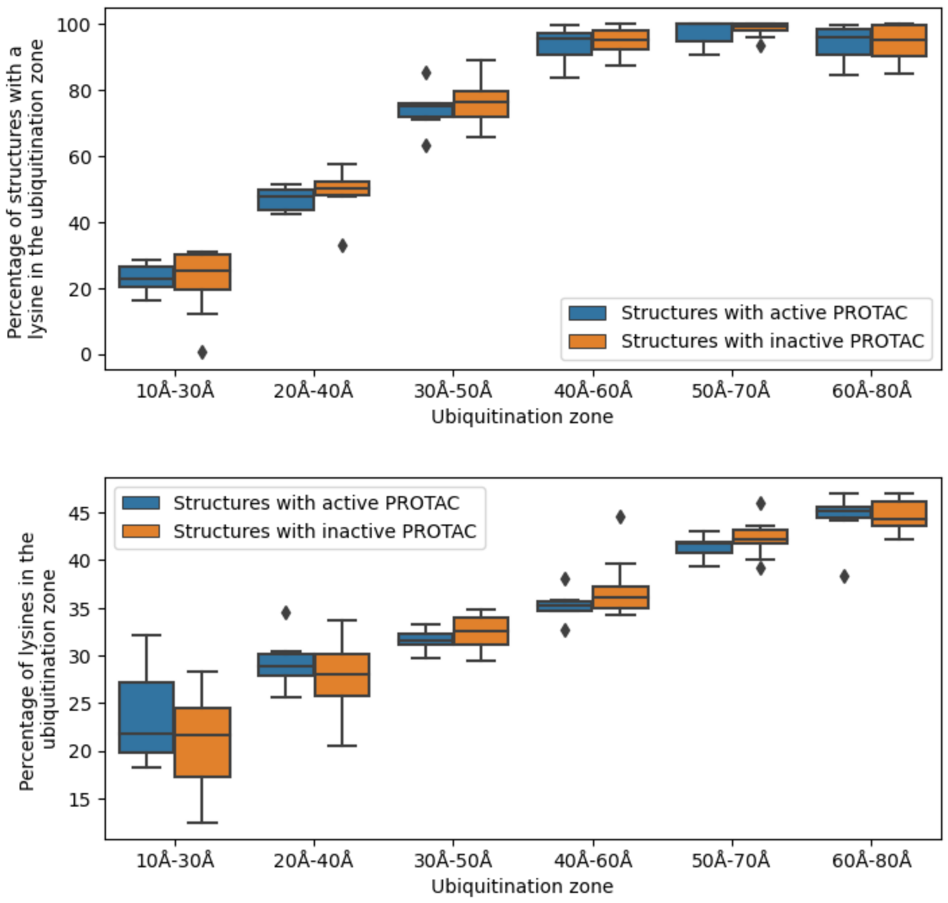

**Figure S2. Different cut-off for ubiquitination zone in the VHL-RING complex.** A) Percentage of structures that have at least one lysine in the specified ubiquitination zone. B) Percentage of lysines in the specified ubiquitination zone.

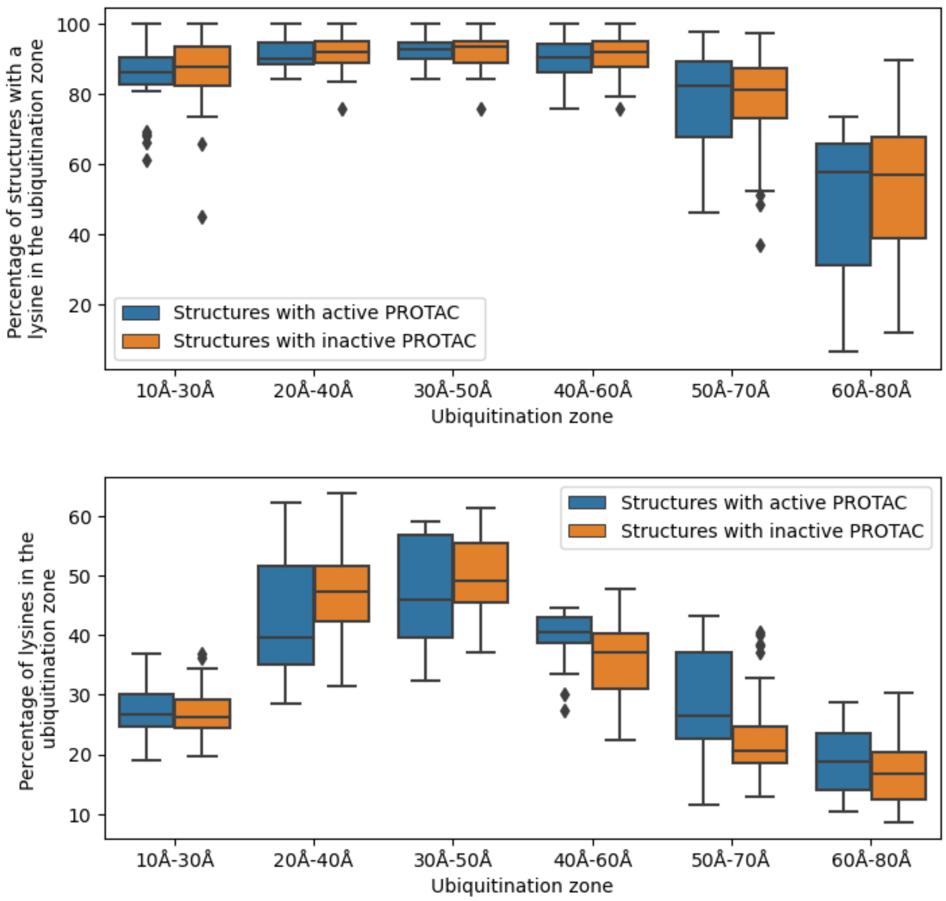

**Figure S3. Different cut-off for ubiquitination zone in the CRBN-RING complex.** A) Percentage of structures that have at least one lysine in the specified ubiquitination zone. B) Percentage of lysines in the specified ubiquitination zone.

**Table S1: Residues selected near chemical handle for HADDOCK docking protocol.**

| Complex | PDB | Residues |
|---|---|---|
| **cIAP-BTK** | **6W74** | 300,301,310,314,315,317,318,328,329 |
| | **3GEN** | 407,484,525 |
| **CRBN-BRD4BD1** | **4TZ4** | 353,355,374,376,377,397 |
| | **3MXF** | 78,79,81,90,91,93,95,138,141,142,145,148 |
| **VHL-BCL** | **4B9K** | 66,67,69,70,77,92,97,99,107,108,110,113 |
| | **3ZLO** | 96,100,101,104,105,106,110,111,132,133,194,195 |
| **VHL-BRD4BD1** | **4B9K** | 66,67,69,70,77,92,97,99,107,108,110,113 |
| | **3MXF** | 78,79,81,90,91,93,95,138,141,142,145,148 |
| **VHL-BRD4BD2** | **4B9K** | 66,67,69,70,77,92,97,99,107,108,110,113 |
| | **5UEO** | 371,374,383,384,388,431,434,436,438 |
| **VHL-FAK1** | **4B9K** | 66,67,69,70,77,92,97,99,107,108,110,113 |
| | **4BRX** | 426,427,430,432,503,504,569 |
| **VHL-SMARCA2** | **5NVX** | 66,67,69,70,77,92,97,99,107,108,110,113 |
| | **6HAZ** | 1407,1416,1417,1419,1462,1465,1469 |
| **VHL-SMARCA4** | **5NVX** | 66,67,69,70,77,92,97,99,107,108,110,113 |
| | **6ZS2** | 1483,1487,1492,1493,1495,1538,1541,1545 |
| **VHL-WDR5/ VHL-WDR5_DC** | **4B9K** | 66,67,69,70,77,92,97,99,107,108,110,113 |
| | **4QL1** | 131,149,258,259 |

**Table S2: Ternary complex prediction input for each complex including the chemical handle definitions and the lowest achieved Ca-RMSD score for each method compared to the crystal structure of the ternary complex**

| Complex | PROTAC | Chemical handle | ICM (Å) | MOE (Å) | PRosettaC (Å) |
|---|---|---|---|---|---|
| cIAP-BTK | 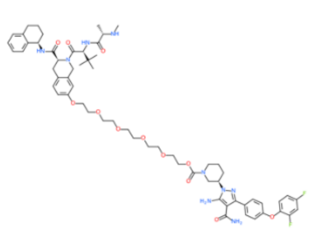 | 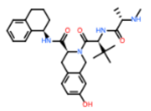<br>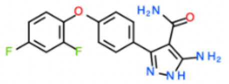 | 10.05 | 21.83 | ND* |
| CRBN-BRD4BD1 | 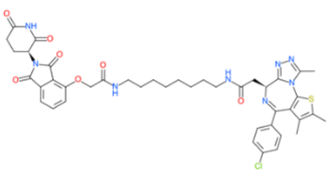 | 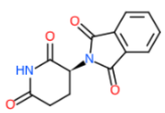<br>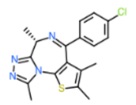 | 5.73 | 3.64 | 3.92 |
| VHL-BCL | 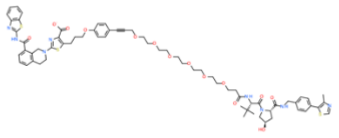 | 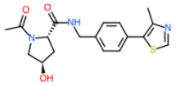<br>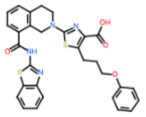 | 2.27 | 14.14 | ND* |
| VHL-BRD4BD1 | 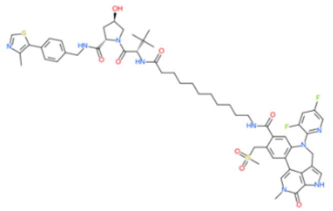 | 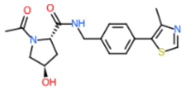<br>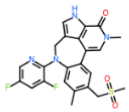 | 1.69 | 4.29 | 7.78 |
| VHL-BRD4BD2 | 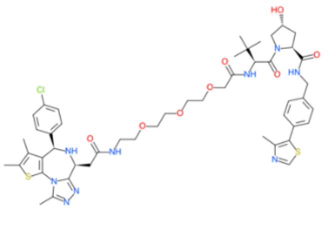 | 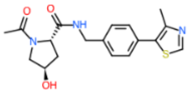<br>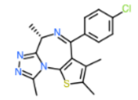 | 6.74 | 3.70 | 6.09 |
| VHL-FAK1 | | 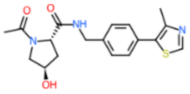 | 4.75 | 9.98 | 11.75 |

| | | | | | |
|---|---|---|---|---|---|
| | 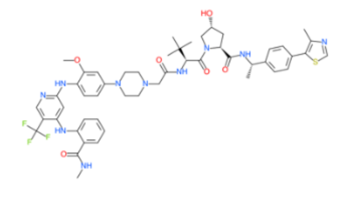 | 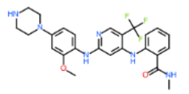 | | | |
| **VHL-SMARCA2** | 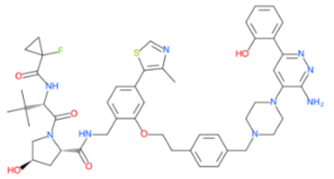 | 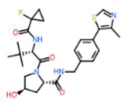<br>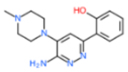 | **3.35** | **12.71** | **16.44** |
| **VHL-SMARCA4** | 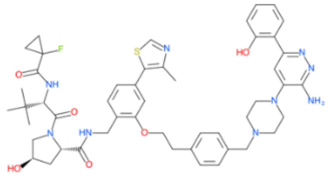 | 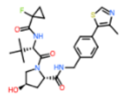<br>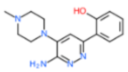 | **3.40** | **5.17** | **15.25** |
| **VHL-WDR5** | 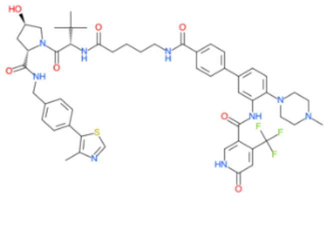 | 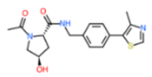<br>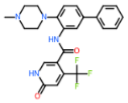 | **7.65** | **10.34** | **18.83** |
| **VHL-WDR5_DC** | 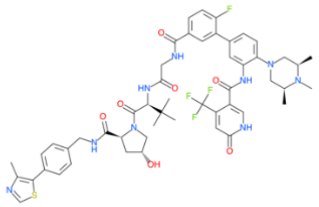 | 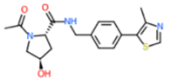<br>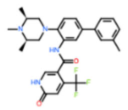 | **3.50** | **8.47** | **11.11** |

*** ND = Not determined: The running time required for the simulation to finish was greater than running time and resources available.**

**Table S3: PROTAC virtual screening input chemical handle definitions**

| Complex | E3 ligase | E3 ligase |
| --- | --- | --- |
| **CRBN-BRAF*** | 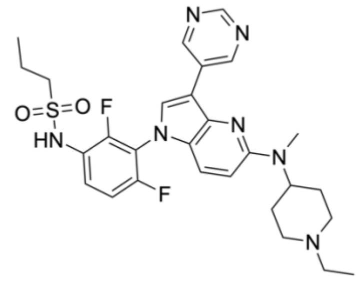 | 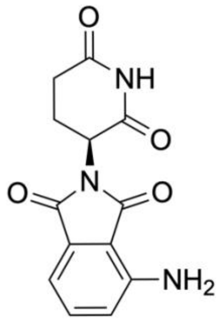 |
| **CRBN-CDK2**** | A:<br>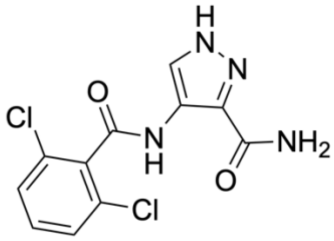<br>F:<br>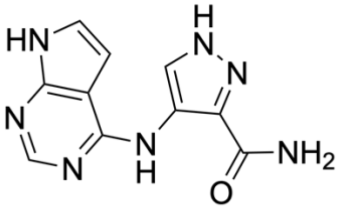 | **1:**<br>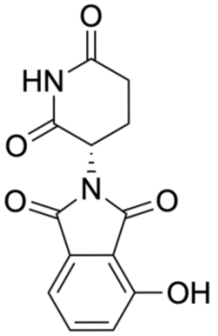<br>2:<br>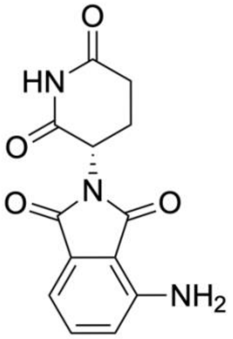 |
| **CRBN-CDK9**** | A:<br>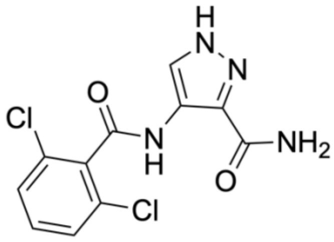 | **1:**<br>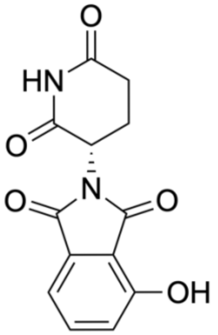 |

| | | |
|---|---|---|
| | F:<br>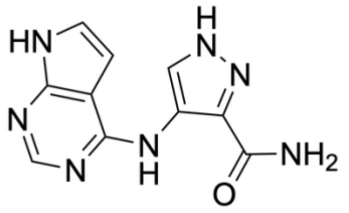 | 2:<br>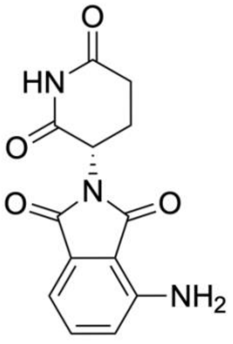 |
| **CRBN-PARP1** | 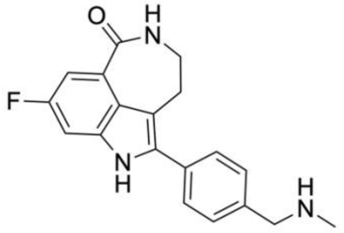 | 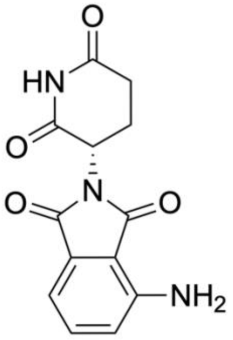 |
| **VHL-WDR5** | **1:**<br>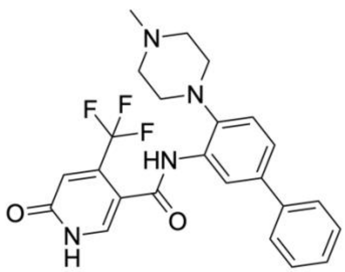<br>2:<br>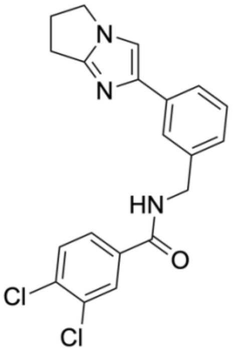 | 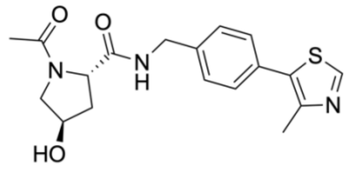 |

* Chemical handle for ICM:

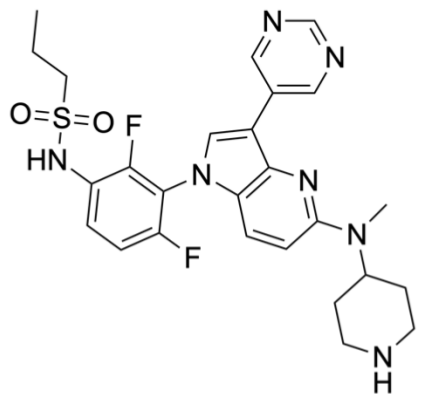

** Chemical handle for PRosettaC:

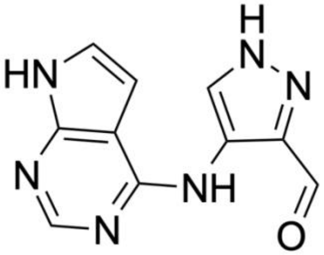

**Table S4: PROTAC virtual screening for CRBN-BRAF including the linker definitions and the ranking for each method.**

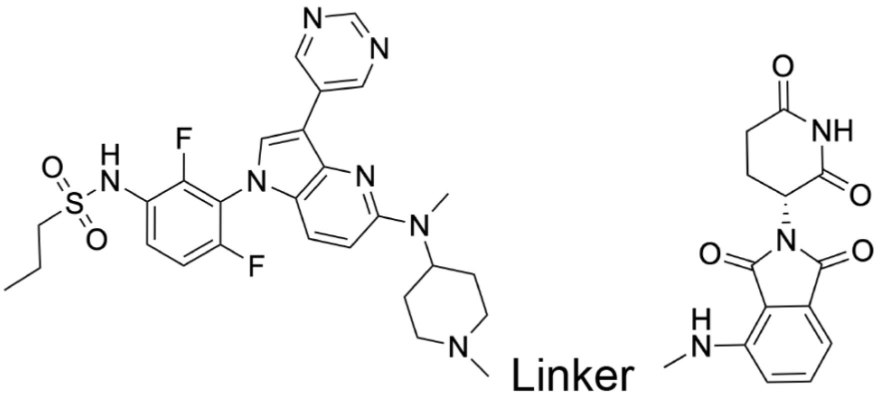

| # | Name | Linker | ICM (ranking) | MOE (ranking) | PRosettaC (ranking) | Activity |
|---|---|---|---|---|---|---|
| 1 | Pmd-1C-PEG3-1C-2C-BI | 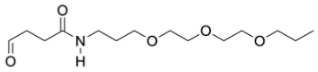 | 5 | 6 | 1 | active |
| 2 | Pmd-C1-Ph-C1-PPZ-C9-BI | 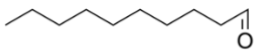 | 3 | 2 | 4 | inactive |
| 3 | Pmd-DODA-C1-BI | 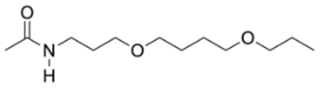 | 2 | 3 | 3 | active |
| 4 | Pmd-PEG1-Trz-PEG2-BI | 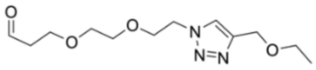 | 6 | 4 | 8 | active |
| 5 | Pmd-PEG3-Az-C1-BI | 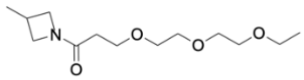 | 8 | 7 | 5 | inactive |
| 6 | Pmd-PEG6-BI | 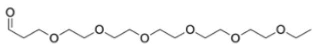 | 1 | 8 | 6 | active |
| 7 | Pmd-PEG8-BI | 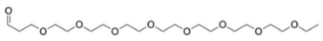 | 7 | 1 | 7 | active |

| | | | | | | |
|---|---|---|---|---|---|---|
| 8 | Pmd-PPZ-C9-BI | 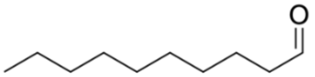 | 4 | 5 | 2 | inactive |

**Table S5: PROTAC virtual screening for CRBN-CDK2 including the linker definitions and the ranking for each method.**

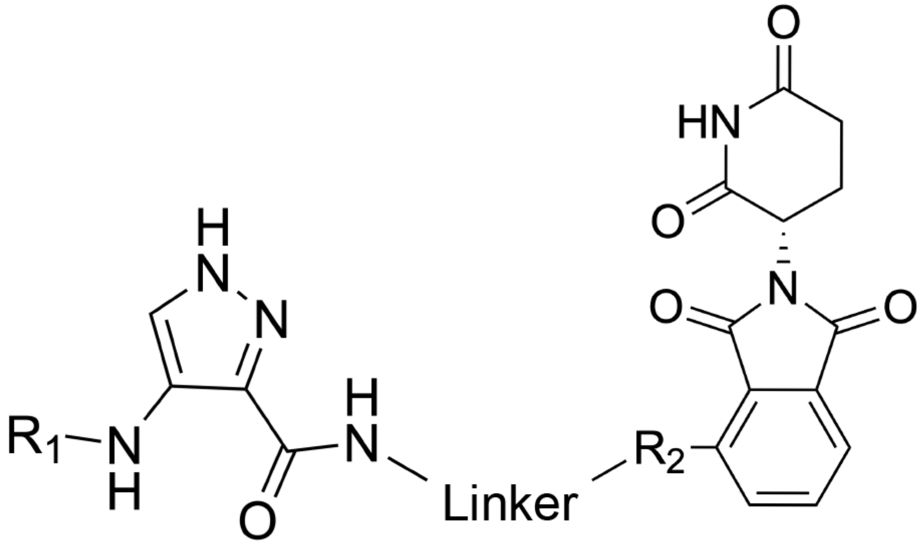

| # | Name | Linker | R1 | R2 | ICM (ranking) | MOE (ranking) | PRosettaC (ranking) | Activity |
|---|---|---|---|---|---|---|---|---|
| 1 | A1 | 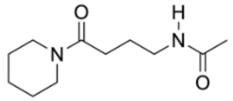 | 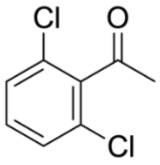 | 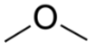 | 4 | 20 | 19 | inactive |
| 2 | A2 | 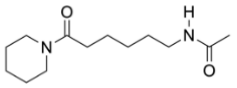 | 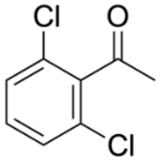 | 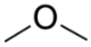 | 17 | 19 | 14 | active |
| 3 | A3 | 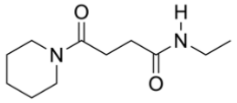 | 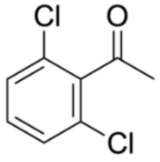 | 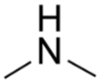 | 11 | 17 | ND* | inactive |
| 4 | A4 | 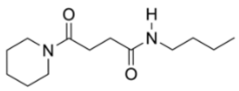 | 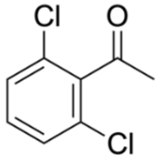 | 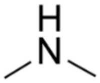 | 8 | 12 | 1 | inactive |

| | | | | | | | | |
|---|---|---|---|---|---|---|---|---|
| 5 | A5 | 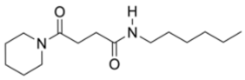 | 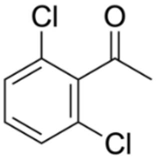 | 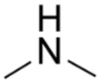 | 1 | 9 | 2 | inactive |
| 6 | A6 | 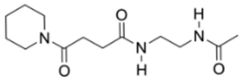 | 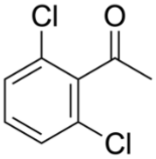 | 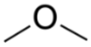 | 6 | 13 | 9 | inactive |
| 7 | A7 | 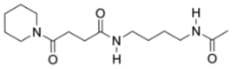 | 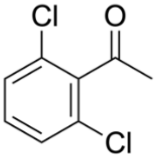 | 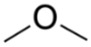 | 10 | 16 | 6 | inactive |
| 8 | A8 | 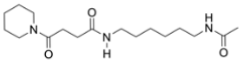 | 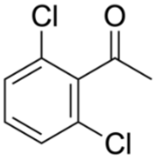 | 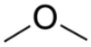 | 3 | 10 | 15 | inactive |
| 9 | A9 | 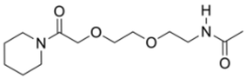 | 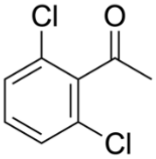 | 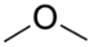 | 2 | 18 | 16 | inactive |
| 10 | A10 | 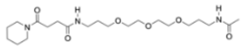 | 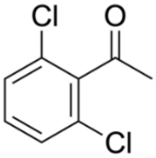 | 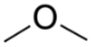 | 18 | 14 | 17 | inactive |
| 11 | F1 | 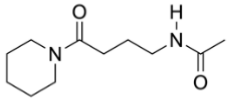 | 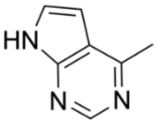 | 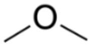 | 5 | 1 | 12 | active |
| 12 | F2 | 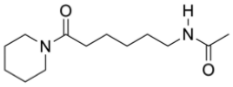 | 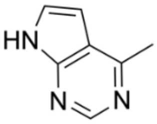 | 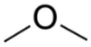 | 14 | 3 | 7 | active |

| | | | | | | | | |
|---|---|---|---|---|---|---|---|---|
| 13 | F3 | 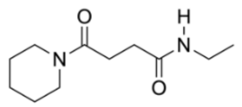 | 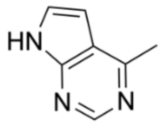 | 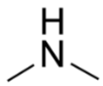 | 13 | 11 | 2 | active |
| 14 | F4 | 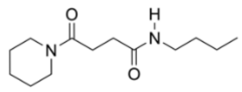 | 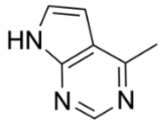 | 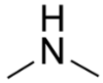 | 15 | 8 | 4 | active |
| 15 | F5 | 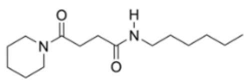 | 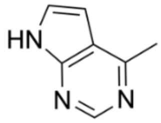 | 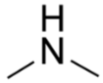 | 9 | 6 | 13 | inactive |
| 16 | F6 | 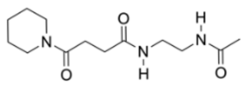 | 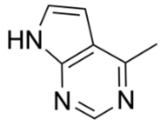 | 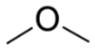 | 7 | 2 | 11 | inactive |
| 17 | F7 | 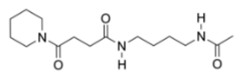 | 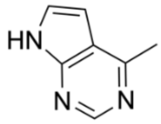 | 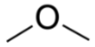 | 12 | 5 | 10 | inactive |
| 18 | F8 | 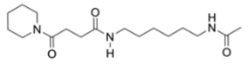 | 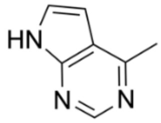 | 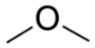 | 20 | 7 | 5 | inactive |
| 19 | F9 | 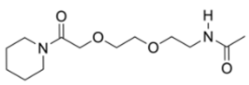 | 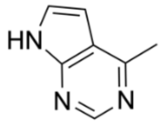 | 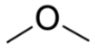 | 16 | 4 | 8 | inactive |
| 20 | F10 | 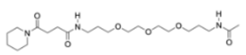 | 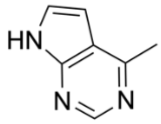 | 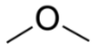 | 19 | 15 | 18 | inactive |

* ND = Not determined: No solutions after running screening.

**Table S6: PROTAC virtual screening for CRBN-CDK9 including the linker definitions and the ranking for each method.**

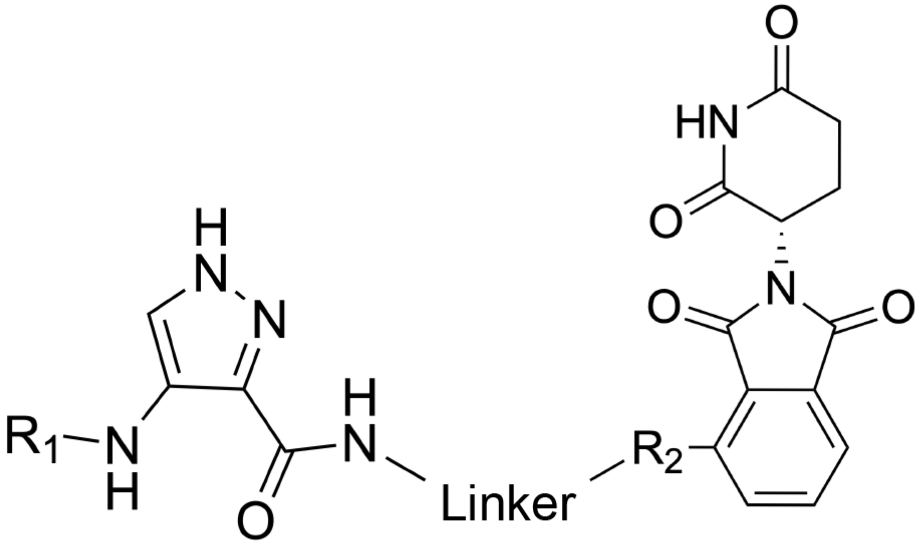

| # | Name | Linker | R1 | R2 | ICM (ranking) | MOE (ranking) | PRosettaC (ranking) | Activity |
|---|---|---|---|---|---|---|---|---|
| 1 | A1 | 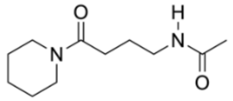 | 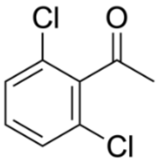 | 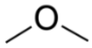 | 18 | 20 | 6 | inactive |
| 2 | A2 | 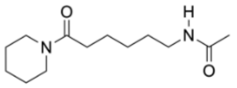 | 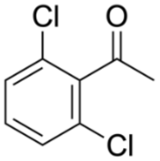 | 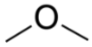 | 8 | 18 | 5 | active |
| 3 | A3 | 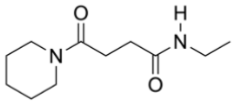 | 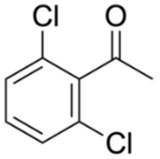 | 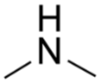 | 13 | 16 | 15 | inactive |
| 4 | A4 | 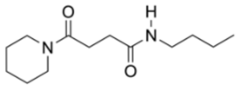 | 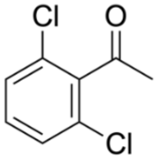 | 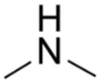 | 3 | 12 | ND* | inactive |

| | | | | | | | | |
|---|---|---|---|---|---|---|---|---|
| 5 | A5 | 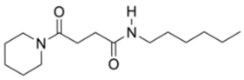 | 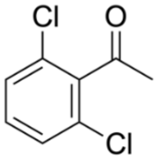 | 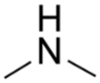 | 11 | 10 | 12 | inactive |
| 6 | A6 | 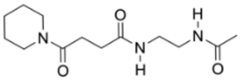 | 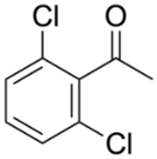 | 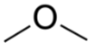 | 6 | 17 | 2 | inactive |
| 7 | A7 | 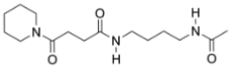 | 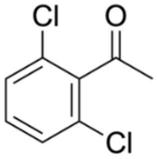 | 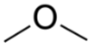 | 15 | 11 | 14 | inactive |
| 8 | A8 | 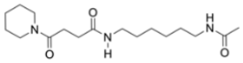 | 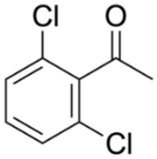 | 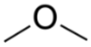 | 16 | 14 | 11 | inactive |
| 9 | A9 | 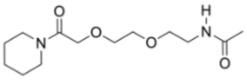 | 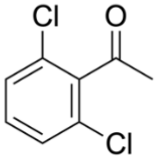 | 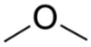 | 19 | 13 | 4 | inactive |
| 10 | A10 | 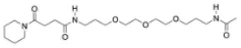 | 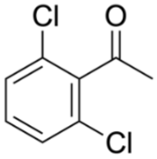 | 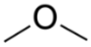 | 20 | 19 | ND* | inactive |
| 11 | F1 | 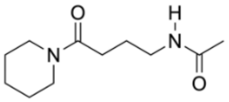 | 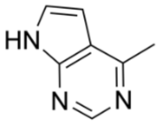 | 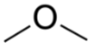 | 5 | 8 | 3 | active |
| 12 | F2 | 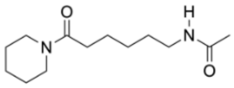 | 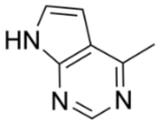 | 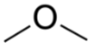 | 4 | 6 | 8 | active |

| | | | | | | | | |
|---|---|---|---|---|---|---|---|---|
| 13 | F3 | 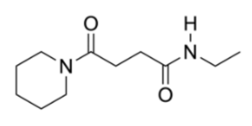 | 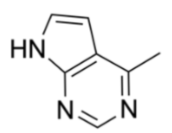 | 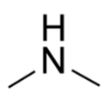 | 1 | 2 | 1 | active |
| 14 | F4 | 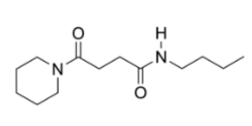 | 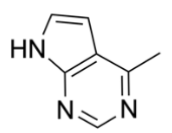 | 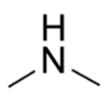 | 2 | 3 | 13 | active |
| 15 | F5 | 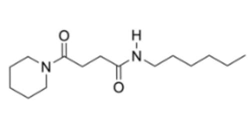 | 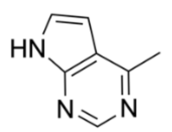 | 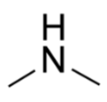 | 7 | 1 | 17 | active |
| 16 | F6 | 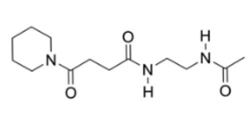 | 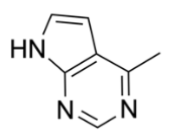 | 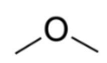 | 9 | 4 | 10 | active |
| 17 | F7 | 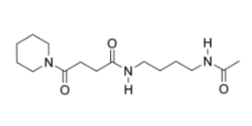 | 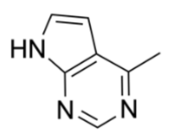 | 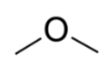 | 12 | 5 | 18 | inactive |
| 18 | F8 | 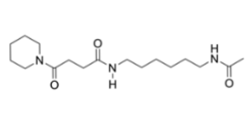 | 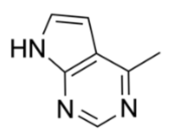 | 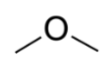 | 14 | 9 | 16 | inactive |
| 19 | F9 | 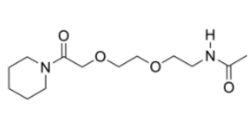 | 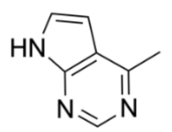 | 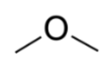 | 10 | 7 | 9 | active |
| 20 | F10 | 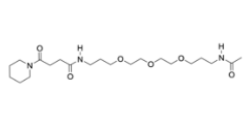 | 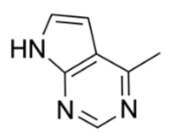 | 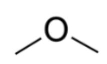 | 17 | 15 | 7 | inactive |

* ND = Not determined: No solutions after running screening.

**Table S7: PROTAC virtual screening input for CRBN-PARP1 including the linker definitions and the ranking for each method.**

A

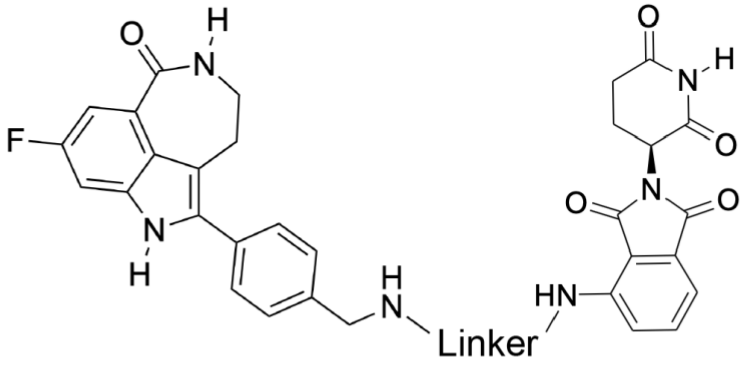

B

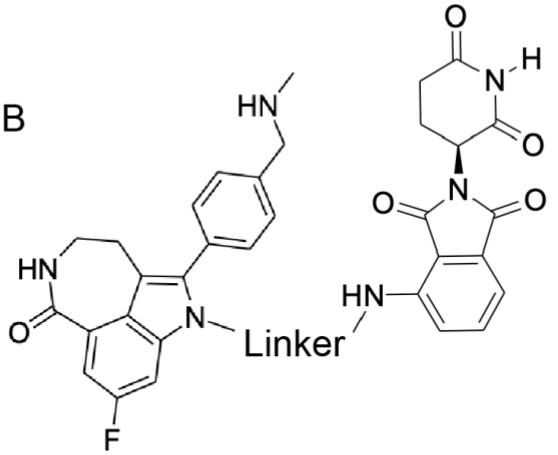

| # | Name | Linker | Attachment point | ICM (ranking) | MOE (ranking) | PRosettaC (ranking) | Activity |
|---|---|---|---|---|---|---|---|
| 1 | iRucaparib-TP1 | 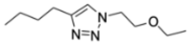 | A | 5 | 1 | 2 | inactive |
| 2 | iRucaparib-TP2 | 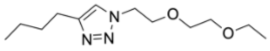 | A | 6 | 12 | 3 | inactive |
| 3 | iRucaparib-TP3 | 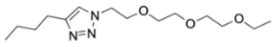 | A | 1 | 3 | 10 | active |
| 4 | iRucaparib-TP4 | 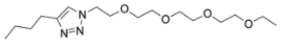 | A | 7 | 13 | 6 | active |
| 5 | iRucaparib-TP5 | 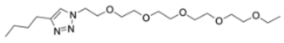 | A | 11 | 11 | 12 | inactive |
| 6 | iRucaparib-AMP3-1 | 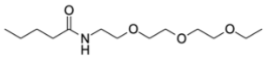 | A | 2 | 5 | 4 | inactive |
| 7 | iRucaparib-AMP3-2 | 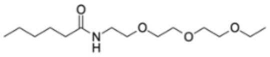 | A | 4 | 8 | 13 | inactive |

| | | | | | | | |
|---|---|---|---|---|---|---|---|
| 8 | iRucaparib-ITP3 | 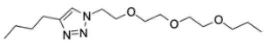 | B | 13 | 4 | 7 | inactive |
| 9 | iRucaparib-ITP4 | 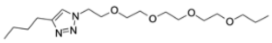 | B | 10 | 6 | 11 | inactive |
| 10 | iRucaparib-AP4 | 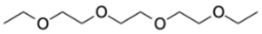 | A | 3 | 2 | 1 | inactive |
| 11 | iRucaparib-AP5 | 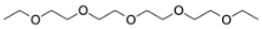 | A | 8 | 10 | 8 | active |
| 12 | iRucaparib-AP6 | 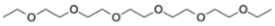 | A | 12 | 9 | 5 | active |
| 13 | iRucaparib-AP7 | 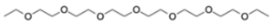 | A | 9 | 7 | 9 | active |

**Table S8: PROTAC virtual screening input for VHL-WDR5 including the linker definitions and the ranking for each method.**

A

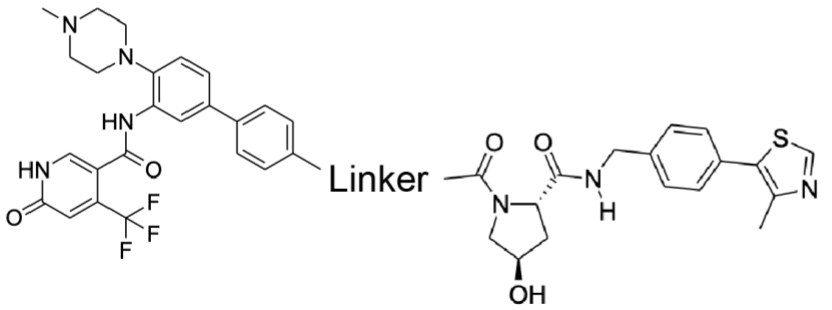

B

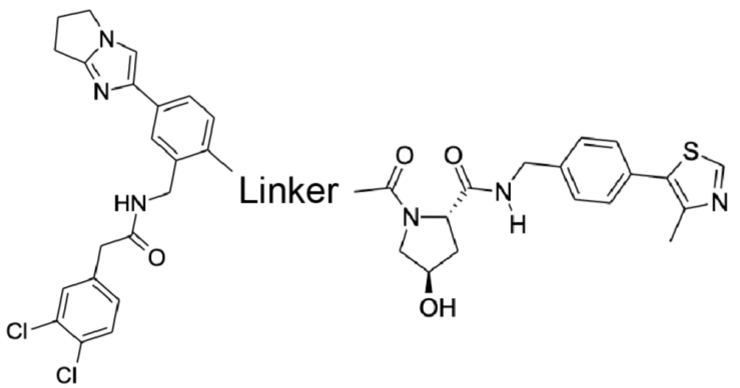

| # | Name | Linker | Chemical handle WDR5 | ICM (ranking) | MOE (ranking) | PRosettaC (ranking) | Activity |
|---|---|---|---|---|---|---|---|
| 1 | 8a | 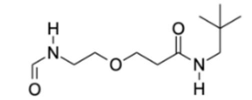 | A | 3 | 9 | 3 | inactive |
| 2 | 8b | 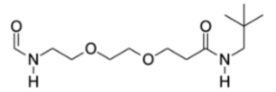 | A | 6 | 8 | 9 | inactive |
| 3 | 8c | 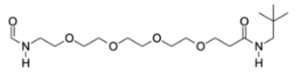 | A | 14 | 7 | ND* | inactive |
| 4 | 8d | 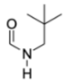 | A | 16 | 16 | ND** | inactive |
| 5 | 8e | 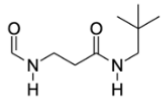 | A | 12 | 4 | 11 | active |

| | | | | | | | |
|---|---|---|---|---|---|---|---|
| 6 | 8f | 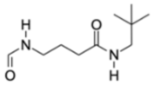 | A | 9 | 6 | 4 | active |
| 7 | 8g | 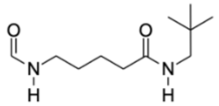 | A | 4 | 3 | 5 | active |
| 8 | 8h | 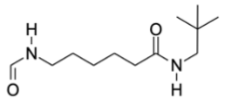 | A | 2 | 2 | 2 | active |
| 9 | 8i | 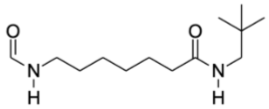 | A | 1 | 1 | 12 | active |
| 10 | 17a | 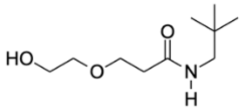 | B | 15 | 10 | 14 | inactive |
| 11 | 17b | 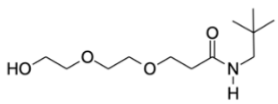 | B | 13 | 11 | 13 | active |
| 12 | 17c | 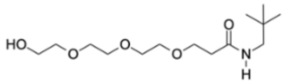 | B | 10 | 14 | 1 | inactive |
| 13 | 17d | 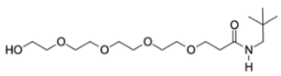 | B | 5 | 15 | 8 | inactive |
| 14 | 17e | 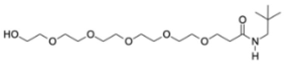 | B | 7 | 5 | 6 | inactive |
| 15 | 17f | 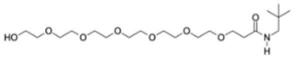 | B | 11 | 12 | 10 | inactive |
| 16 | 17g | 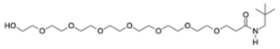 | B | 8 | 13 | 7 | inactive |

* ND = Not determined: The running time required for the simulation to finish was greater than running time and resources available.

** ND = Not determined: No solutions after running screening.

**Table S9. Common ternary conformations induced by active PROTACs.** The complex is presented with the E3 ligase in grey and target in color. The colors represent the different common conformations observed between active PROTACs and not observed among inactive PROTACs.

| Complex | Color conformation | Number of PROTACs that induce conformation | PROTACs that induce conformation |
|---|---|---|---|
| CRBN-BRAF | 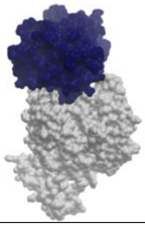 | 2 | ID: 1,4<br><br>1) Pmd-1C-PEG3-1C-2C-BI<br>4) Pmd-PEG1-Trz-PEG2-BI |
| CRBN-BRAF | 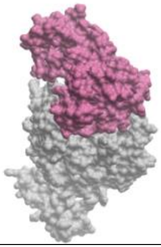 | 2 | ID: 1,3<br><br>1) Pmd-1C-PEG3-1C-2C-BI<br>3) Pmd-DODA-C1-BI |
| CRBN-BRAF | 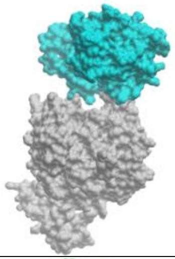 | 3 | ID: 3,4,7<br><br>3) Pmd-DODA-C1-BI<br>4) Pmd-PEG1-Trz-PEG2-BI<br>7) Pmd-PEG8-BI |
| CRBN-BRAF | 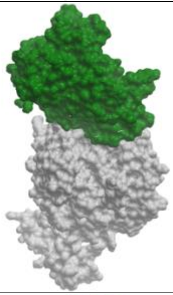 | 2 | ID: 3,6<br><br>3) Pmd-DODA-C1-BI<br>6) Pmd-PEG6-BI |
| CRBN-BRAF | 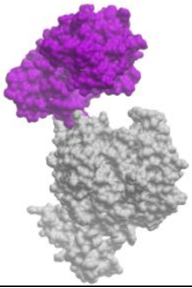 | 2 | ID: 3,4<br><br>3) Pmd-DODA-C1-BI<br>4) Pmd-PEG1-Trz-PEG2-BI |

| | | | |
|---|---|---|---|
| CRBN-CDK2 | 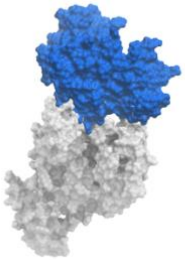 | 2 | ID: 2,12<br><br>2) A2<br>12) F2 |
| CRBN-CDK9 | 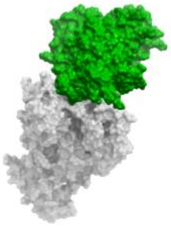 | 2 | ID: 11,14<br><br>11) F1<br>14) F4 |
| CRBN-CDK9 | 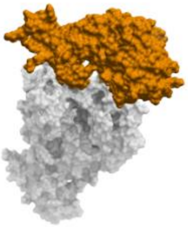 | 2 | ID: 12,16<br><br>12) F2<br>16) F6 |
| CRBN-CDK9 | 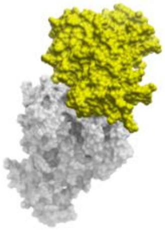 | 2 | ID: 13,15<br><br>13) F3<br>15) F5 |
| CRBN-PARP1 | 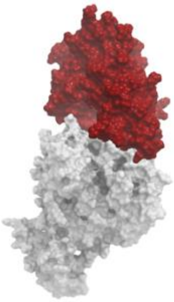 | 2 | ID: 4,13<br><br>4) iRucaparib-TP4<br>13) iRucaparib-AP7 |
| VHL-WDR5 | 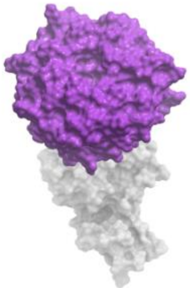 | 2 | ID: 5,7<br><br>5) 8e<br>7) 8g |

| | | | |
|---|---|---|---|
| VHL-WDR5 | 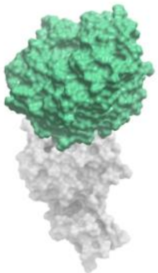 | 2 | ID: 5,6<br><br>5) 8e<br>7) 8g |
| VHL-WDR5 | 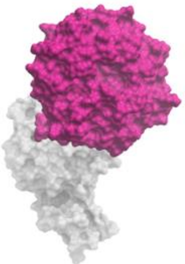 | 3 | ID: 6,7,8<br><br>6) 8f<br>7) 8g<br>8) 8h |
| VHL-WDR5 | 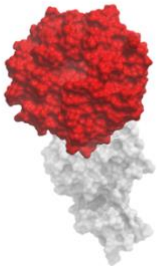 | 2 | ID: 8,9<br><br>8) 8h<br>9) 8i |
